# RNA polymerase clamp movement aids dissociation from DNA but is not required for RNA release at intrinsic terminators

**DOI:** 10.1101/453969

**Authors:** Michael J Bellecourt, Ananya Ray-Soni, Alex Harwig, Rachel Anne Mooney, Robert Landick

## Abstract

In bacteria, disassembly of elongating transcription complexes (ECs) can occur at intrinsic terminators in a 2-3 nucleotide window after transcription of multiple kilobase pairs of DNA. Intrinsic terminators trigger pausing on weak RNA-DNA hybrids followed by formation of a strong, GC-rich stem-loop in the RNA exit channel of RNA polymerase (RNAP), inactivating nucleotide addition and inducing dissociation of RNA and RNAP from DNA. Although the movements of RNA and DNA during intrinsic termination have been studied extensively leading to multiple models, the effects of RNAP conformational changes remain less well-defined. RNAP contains a clamp domain that closes around the nucleic-acid scaffold during transcription initiation and can be displaced by either swiveling or opening motions. Clamp opening is proposed to promote termination by releasing RNAP-nucleic acid contacts. We developed a cysteine-crosslinking assay to constrain clamp movements and study effects on intrinsic termination. We found that biasing the clamp into different conformations perturbed termination efficiency, but that perturbations were due primarily to changes in elongation rate, not the competing rate at which ECs commit to termination. After commitment, however, inhibiting clamp movements slowed release of DNA but not of RNA from the EC. We also found that restricting trigger-loop movements with the RNAP inhibitor microcin J25 prior to commitment inhibits termination, in agreement with a recently proposed multistate-multipath model of intrinsic termination. Together our results support views that termination commitment and DNA release are separate steps and that RNAP may remain associated with DNA after termination.

**Highlights:**

- Disulfide bond crosslinks probe the role of the RNAP clamp domain in termination
- RNA but not DNA can release at terminators when the RNAP clamp is closed
- Restricting RNAP clamp movement affects elongation rate more than termination rate
- Inhibiting TL conformational flexibility impairs both RNA and DNA release

## INTRODUCTION

Termination of transcription by RNA polymerase (RNAP) is the essential process in all organisms that must occur at the end of every functional transcription unit (*i.e*., a gene or polycistronic operon) to release the newly synthesized RNA product and dissociate RNAP from the DNA. Failure to terminate efficiently has deleterious effects on the cell, including but not limited to unregulated expression of downstream genes [1] and collisions with replication complexes that can cause double-stranded breaks in genomic DNA [2]. Bacteria have evolved two mechanisms to promote transcription termination at the ends of transcription units. Rho-dependent termination relies upon the RNA helicase Rho to disassemble the elongation complex (EC), whereas intrinsic termination induces EC disassembly *via* a signal encoded by the DNA and RNA comprised of a palindromic GC-rich dyad immediately followed by an ~8-nt U-rich tract [3].

As described by thermodynamic models for intrinsic termination, termination efficiency (TE) is determined by the relative free energy barriers to termination *vs.* elongation at each template position [4, 5]. At most positions, the EC is strongly biased towards continued elongation due to rapid nucleotide addition (up to 100 s^-1^) and the high stability of the EC resulting from combined van der Waals and polar interactions between RNAP and the nucleic-acid (NA) scaffold. The NA scaffold is comprised of a 9–10-bp RNA–DNA hybrid within a ~12-bp transcription bubble, ~5 nt of single-stranded RNA threading out of the RNA exit channel, and ~18 bp of double-stranded downstream DNA [6–9].

Intrinsic terminators facilitate entry into the termination pathway through the U-tract and its surrounding sequences, which promote elemental pausing of the EC [10–13] (Figure 1A). The U-tract pause slows elongation and provides time for the GC-dyad to begin folding into a terminator hairpin (T_hp_), possibly inducing a transient hairpin-stabilized pause [14]. Unlike conventional hairpin pause signals, the juxtaposition of the weak rU:dA hybrid [15] and additional strong rG:rC base pairs at a terminator allow extension of the hairpin by an additional 2–3 bp, coming to within 8 nt of the RNA 3′ end. This process, called T_hp_ completion, appears to be rate-limiting for termination because it creates steric clashes in the EC that can only be resolved by partial melting of the RNA-DNA hybrid [6]. At terminators like *t*_*hisL*_ and λ*t*_R2_ with perfect or near-perfect U-tracts (Figure 1B), this melting is achieved by shearing of the hybrid that pulls 2–3 conserved 5′ Us of the U-tract into the exit channel, permitting T_hp_ completion [6, 16] and EC destabilization [17]. Shearing is thermodynamically unfavorable at terminators like *t*_500_ with “imperfect” U-tracts interrupted by at least one GC bp (Figure 1B). Instead, the T_hp_ exerts a pushing force on RNAP, driving forward translocation and shortening of the hybrid without nucleotide addition due to stronger base-pairing of the RNA with template DNA [16, 18]. Regardless of the path to hybrid shortening, after T_hp_ completion the RNA 3′ end is displaced from the active site, preventing catalysis and committing the EC to dissociation.

**Figure 1.**
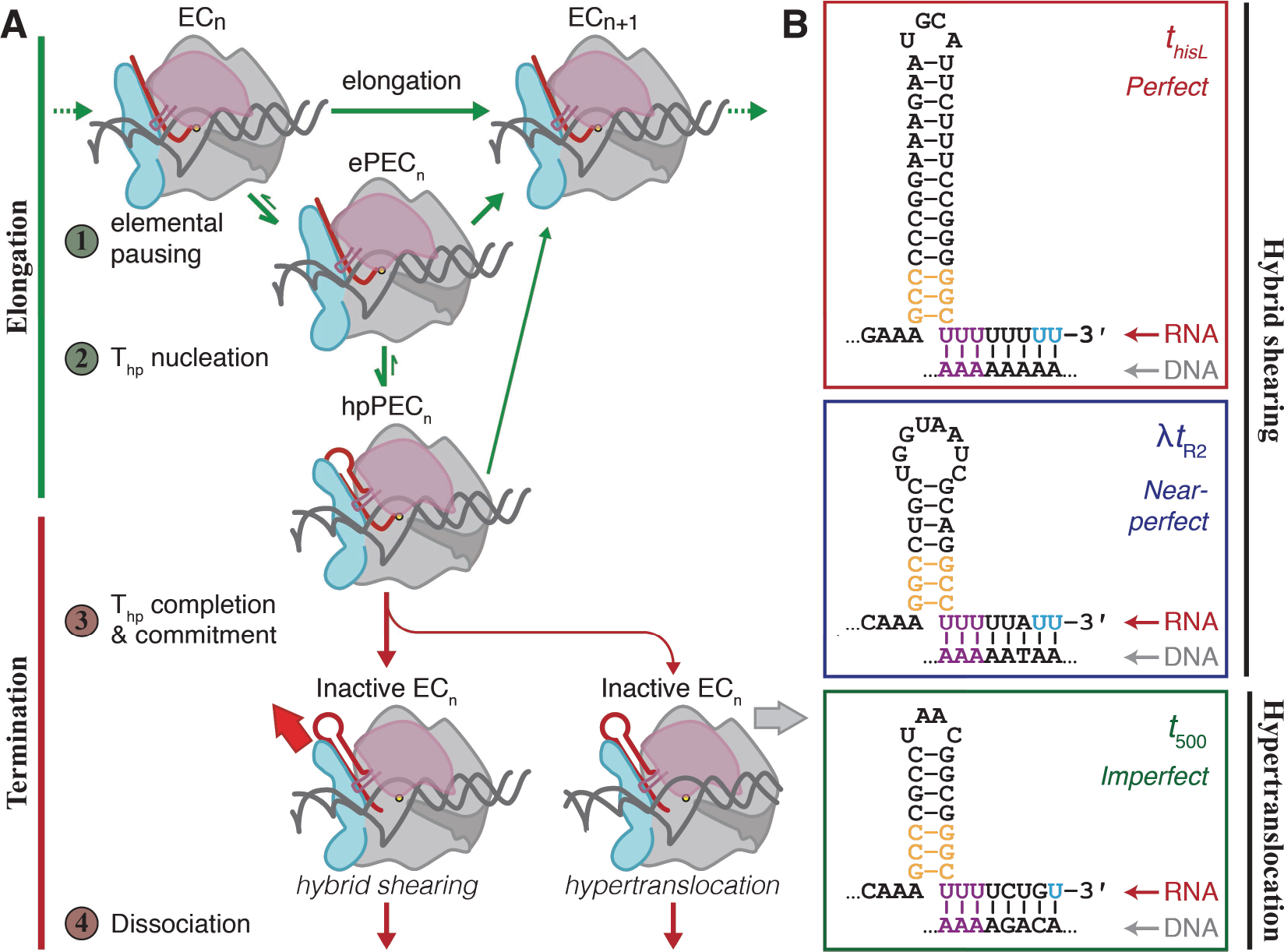
The mechanism of intrinsic termination. (**A**) The four major steps of the intrinsic termination pathway include steps that impact the elongation mechanism (green arrows) and those that impact the termination mechanism (red arrows). Each step along the mechanism is modulated by complex interactions between the nucleic-acid scaffold (DNA, grey; RNA, red) and RNAP itself. ePEC, elemental paused elongation complex; hpPEC, hairpin-stabilized pause elongation complex. (**B**) Three model intrinsic terminators used in these studies. The two major components of an intrinsic terminator include a GC-rich dyad that folds into an RNA hairpin and a U-rich tract immediately downstream. The strength of the U-tract, as measured by its uridine-content, determines its class of terminator. Violet uridine bases indicate the most highly-conserved uridines that melt upon pairing of the final 2-3 G-C bases at the stem of the T_hp_ (orange). Blue uridine bases indicate the sites of termination.

In contrast to the movements of the NA scaffold, the movements of RNAP that contribute to termination are poorly understood. RNAP is composed of several mobile parts whose coordinated movements catalyze the nucleotide addition cycle. Specific conformational changes are proposed to facilitate termination [19–24]. One hypothesis is that opening of the clamp is necessary for termination [22–24, 25]. During initiation the clamp moves from an open to a closed state that persists throughout elongation (Figure 2A) [21–22, 26, 27]. Clamp closure establishes RNAP–NA interactions that stabilize the EC [21, 26] and prevents transcription bubble collapse because double-stranded DNA cannot fit into the narrowed main cleft [22]. Clamp opening at a terminator could therefore facilitate termination by breaking the stabilizing RNAP–NA interactions, [21, 26] enabling bubble collapse and DNA release [19–22, 23, 28, 29], but the extent to which clamp opening is required for termination remains unknown. The *his* paused EC includes a pause RNA hairpin similar to the incomplete T_hp_, and studies suggest this hairpin promotes rotation of the clamp toward the RNA exit channel (orthogonal to clamp opening) as part of a rigid body rotation of a swivel module that includes the clamp and is required for pause stabilization [30]. However, it is unknown if clamp swiveling is required for intrinsic termination.

**Figure 2.**
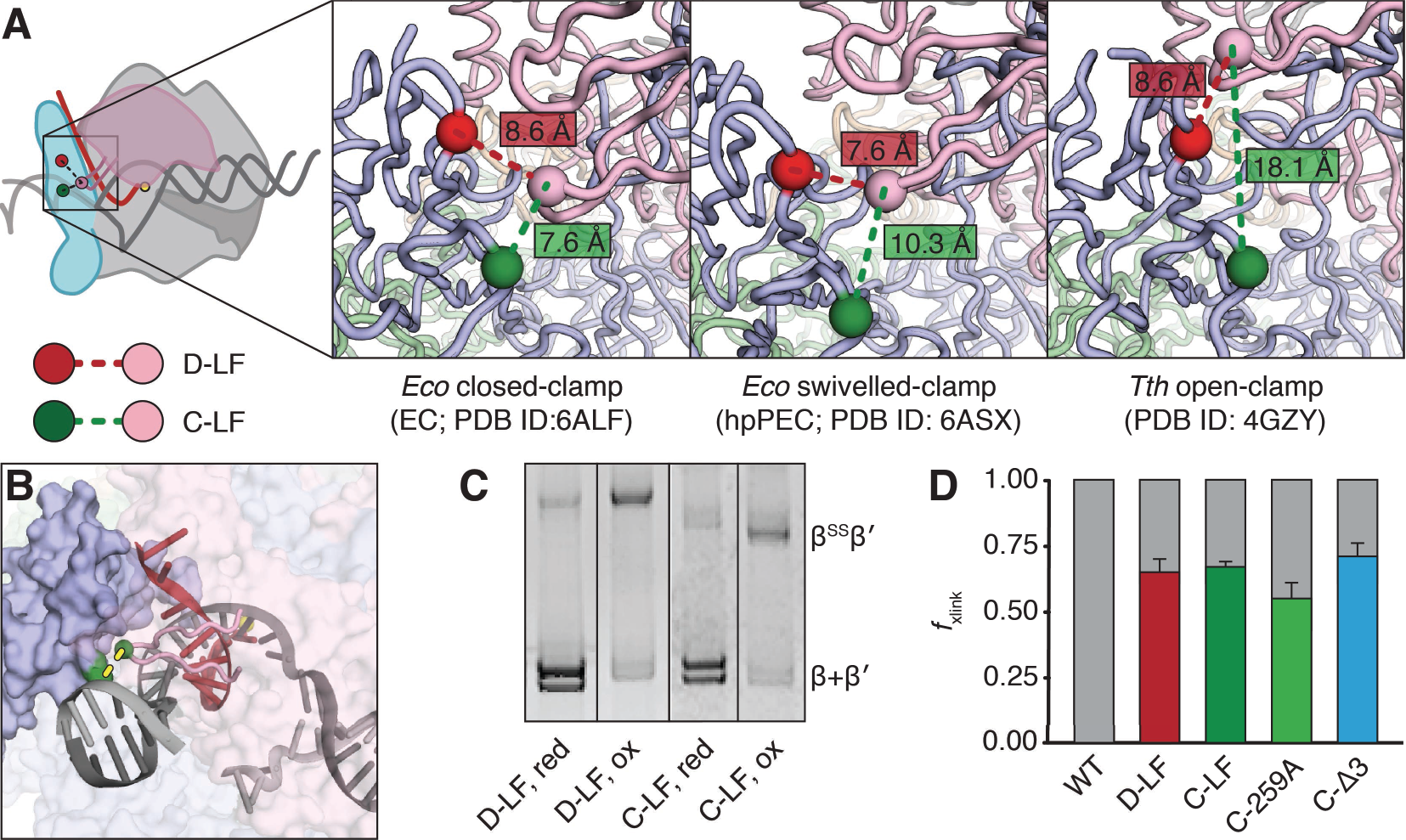
Disulfide crosslinks between the RNAP flap and lid. (**A**) View of cysteine substitutions and insertions within the flap and lid from the same angle across ECs with three distinct clamp conformations. Crosslinking between βC1044 (green sphere) and β′C258 (pink sphere) biases the clamp towards a closed-conformation, whereas crosslinking between βC843 (red sphere) and β′258 biases displacement of the clamp into either the swiveled or open conformation. Panels depict all three cysteines sites for comparison of relative distances between Cα atom but actual RNAP variants have only two cysteines. Closed-clamp EC, PDB ID: **<u>6ALF</u>**[27], swiveled-clamp PEC, PDB ID: **<u>6ASX</u>** [30], open-clamp PEC, PDB ID: **<u>4GZY</u>** [21]. (**B**) An alternative view of the *E. coli* EC (PDB ID: **<u>6ALF</u>**) depicting both protein and NA elements and the relative position of the C-LF flap-lid disulfide, with residues involved colored green with a dashed yellow line. The flap is depicted as a solid blue surface and the lid as a pink ribbon. Other elements of RNAP have been rendered transparent for clarity. The yellow sphere indicates active-site Mg^2+^. (**C**) Representative non-reducing SDS-PAGE gel of D-LF and C-LF under reducing and oxidizing conditions that disfavor and favor disulfide formation, respectively. Crosslink formation causes a supershift of the crosslinked RNAP, allowing for measurement of crosslinking efficiency. Analyses were performed on washed, bead-bound ECs (see Figure 3) rather than bulk RNAP in solution in order to determine crosslinking efficiencies of the transcription-competent complexes. (**D**) Crosslinking efficiencies of the five RNAPs in ECs used to investigate transcriptional elongation and termination commitment on the ligation scaffold. Error bars represent the standard deviation at n > 3.

A second hypothesis is that the T_hp_, once formed, can invade the main channel to contact the folded trigger loop (TL), disrupting RNAP–NA contacts and forcing clamp opening [19]. Other findings suggest that T_hp_–TL contact may reflect post-termination RNAP-RNA binding [17] and that termination may proceed through multiple, alternative RNAP conformational states whose formation is aided by TL flexibility [14]. However, the role of clamp motions during steps in the termination mechanism remains unexplored.

To investigate the role of RNAP clamp and TL motions during intrinsic termination, we developed an experimental approach in which ECs were reconstituted on long DNA templates tethered to paramagnetic beads, followed by treatment to restrict RNAP movements, either by (*i*) oxidation of a pair of cysteine residues engineered into variant RNAPs to suppress clamp movements via disulfide bond formation, or (*ii*) addition of an antimicrobial peptide to block TL movement. Using this experimental approach, we tested the contributions of RNAP conformational freedom to the termination mechanism. We found that clamp conformation principally affects intrinsic termination by affecting elongation rate and that clamp movement, likely opening, is required for DNA but not RNA release at terminators.

## RESULTS

### Restricting clamp movements by Cys-pair crosslinking alters elongation rate

To explore how clamp conformation impacts the intrinsic termination mechanism, we expanded a set of lid-flap cysteine-pair (Cys-pair), variant RNAPs (Figure 2A). The original two RNAP variants, C-LF (closed-clamp, lid-flap; βP1044C β′258iC) and D-LF (displaced-clamp, lid-flap; βT843C β′258iC), were used previously to characterize the role of clamp movements at the *his* pause, a hairpin-stabilized pause encoded in the histidine operon leader region [31]. These Cys-pair crosslinks connect the lid, a flexible β′ module that extrudes from the clamp at the upstream fork junction (Figure 1A) and facilitates separation of the nascent transcript from the template DNA strand [32], to the β flap, which forms one wall of the RNA exit channel. The C-LF crosslink stabilizes the clamp in a closed conformation [26, 27], whereas the D-LF crosslink biases the clamp away from the closed conformation and toward either the open [21] or swiveled conformations [21–30, 33]. Since either clamp opening or swiveling could occur during termination, we designate D-LF as favoring clamp displacement.

The intrinsically mobile nature of the lid means that both the C-LF and D-LF Cys-pairs can bias the clamp to favor closed or displaced positions but do not lock it into these positions. To reduce lid flexibility, we constructed a C-Δ3 Cys-pair RNAP with a three amino-acid deletion directly adjacent to the site of cysteine insertion at the tip of the lid loop. One of the deleted residues in this construct, R259, is proposed to contact the −9 base of the template strand in the post-translocated RNA–DNA hybrid [26, 27]. To distinguish whether any differences in elongation and termination behavior by C-Δ3 were due to the Cys-pair crosslink or the loss of R259–NA contacts, we also constructed a C-259A RNAP (βP1044C β′258iC R259A) that encodes a β′R259A substitution but is otherwise identical to C-LF. The D-LF, C-LF, and C-Δ3 RNAPs all formed Cys-pair crosslinks with ~70% efficiency in ECs reconstituted on nucleic-acid scaffolds after treatment with the oxidizing agent diamide (Figure 2C–D; see also Ref. [31]). C-259A formed crosslinks in 55 ± 6% of the EC (Figure 2D). Because the Cys-pair crosslinking efficiency can vary among conditions, we determined the crosslinking efficiency in each experiment (see figure legends).

To analyze the effects of Cys-pair disulfides on termination, we developed an assay that allows crosslink formation in active ECs and subsequent testing of their effects on different terminators ligated downstream of the ECs (Figure 3A) [34, 35]. RNAP was initially reconstituted on a short oligonucleotide scaffold bearing a template-strand 5′ phosphate and 4-nt StyI overhang for subsequent ligation. Downstream sequences of interest (*e.g.*, a pause signal or a terminator) were generated as a piece of double-stranded DNA by PCR of template plasmids followed by StyI-digestion of a site added in the nontemplate-strand PCR primer. These DNAs were then tethered to paramagnetic streptavidin beads through a 5′ biotin moiety on the template-strand PCR primer. The G17-reconstituted EC and the bead-bound downstream sequences were ligated together, followed by EC extension to the A26 position with ATP and [α-^32^P]GTP. Bead immobilization of the ECs allowed a series of buffer exchanges that washed off RNAPs not formed into active ECs. The radiolabeled A26 ECs were then treated with diamide to generate Cys-pair crosslinks. A sample was taken to determine the fraction of ECs containing disulfide crosslinks. Transcription was then restarted for the termination assay by addition of all four NTPs, and RNAP’s progress along the DNA template was monitored by the removal and quenching of samples at various time points. Under reducing conditions, all ECs resumed transcription from A26 at high efficiencies of >90%. Treatment of the same ECs with diamide to form a Cys-pair crosslink did cause small reductions in restart efficiencies, but these changes were minimal (≥86% ECs still resumed transcription). Since the crosslinking efficiency of A26 ECs was ≥55%, most crosslinked ECs must have resumed transcription.

**Figure 3.**
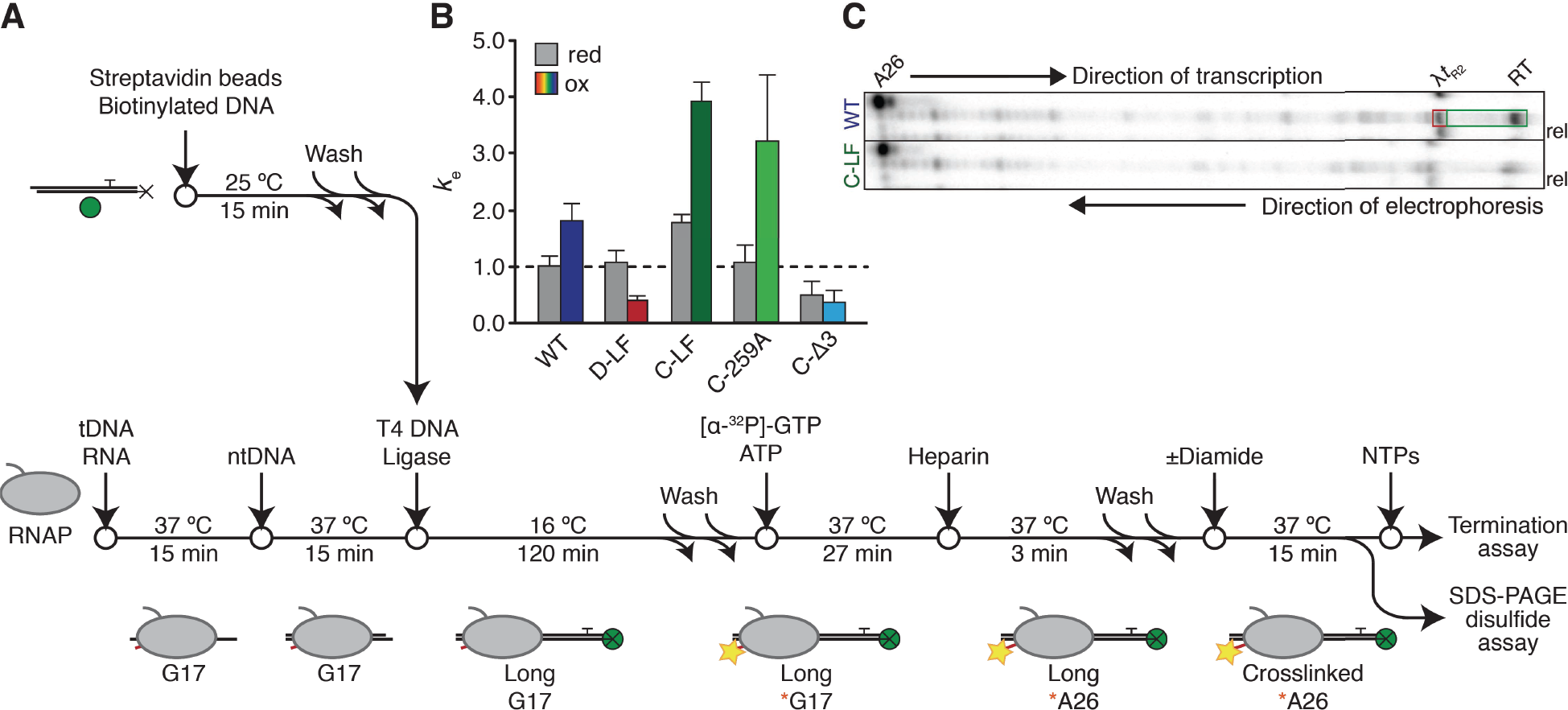
The bead-tethered ligation scaffold assay. (**A**) Assay schematic. Manipulations of the oligonucleotide scaffold and the RNAP are exploited to reduce the homogeneity of complexes. BE, buffer exchange; the magnetic properties of the Streptavidin bead are exploited to eliminate unbound RNAPs and ECs that fail to ligate downstream DNAs of interest. Yellow stars indicate incorporation of radiolabel into RNA (red). Green spheres indicate paramagnetic beads (not to scale); black Xs represent the 5′ biotin moiety on the template strand. (**B**) Elongation properties of Cys-pair RNAPs under reducing (grey bars) and oxidizing (colored bars) conditions. Relative composite pseudo-first-order elongation rate constants are normalized to that of the wild-type EC under reducing conditions to facilitate direct comparisons of elongation rates and subsequent evaluations of termination processes, as indicated by the dashed black line. Error bars indicate standard error as calculated by KinTek Explorer, n = 3. Crosslinking efficiencies under these experimental conditions were: D-LF, 65±5%; C-LF, 67±2%; C-259A, 55±6%; C-A3, 71±5%. (**C**) Representative urea PAG showing *in vitro* transcription of WT and C-LF RNAPs termination at λ*t*_R2_. ECs are radiolabeled and walked to the A26 RNA position. Samples were taken before and after (10 min) addition of NTPs. RT, termination readthrough. Positions of λ*t*_R2_ migration are not demonstrably different between reducing and oxidizing conditions. To demonstrate that termination was occurring, at the end of the reaction mixtures were applied to a magnetic rack and RNAs released from the bead-tethered DNA template were collected. TE is calculated as the fraction of radioactive counts enclosed in the part of the gel indicated by the red box over the sum of radioactive counts in the parts of the gel indicated by the red terminator and green readthrough boxes.

The baseline elongation rate of the non-crosslinked RNAPs varied (under reducing conditions; Figure 3B). To facilitate comparisons, we normalized each elongation rate to that of the wild-type EC under reducing conditions. Under these conditions, which disfavor formation of disulfide crosslinks, both D-LF and C-LF RNAPs elongated more rapidly than the wild-type but to varying degrees (~1.1- and ~1.8-fold faster, respectively). Treatment with diamide to form Cys-pair crosslinks perturbed the elongation rate of all four Cys-pair RNAPs but also affected the wild-type (non-crosslinked) RNAP. Under these oxidizing conditions, wild-type RNAP elongated ~1.8-faster than it did under reducing conditions, suggesting an undefined role for redox state in the elongation behaviors of RNAP. Oxidation of the D-LF RNAP to bias the clamp into the displaced conformation slowed elongation to an average rate only 38% of that of the wild-type RNAP in the reduced state, and 36% relative to its own rate under reducing conditions. C-LF RNAP, on the other hand, extended at a much faster rate relative to both the wild-type baseline (~3.9-fold faster) and itself under reducing conditions (~2.2-fold faster). In contrast, the C-Δ3 EC was impaired for elongation under both oxidizing and reducing conditions. Under reducing conditions, the C-Δ3 RNAP elongated slower by a factor of ~2.0 compared to wild-type. Upon formation of the crosslink, its average elongation rate was further reduced to ~37% that of the baseline wild-type rate, or ~73% of its rate under reducing conditions. C-259A, on the other hand, behaved more similarly to C-LF than to C-Δ3, elongating at nearly the same average rate as the wild-type under reducing conditions, and ~3.2-fold faster upon formation of the crosslink. These results suggest that shortening of the lid loop of a Cys-pair crosslinked RNAP and not the removal of R529 was principally responsible for slow elongation of the C-Δ3 RNAP.

Examination of the transcriptional behavior of these ECs suggested that changes in elongation rate upon disulfide formation were principally due to changes in pause behavior (Figure S1). For instance, formation of the C-LF crosslink appeared to eliminate most weaker pauses along the DNA template and, in agreement with previous findings [30, 31], reduced the strength of the *his* pause. On the other hand, formation of the D-LF disulfide extended the efficiency and duration of weak pauses and caused the EC to dwell at the *his* pause for tens of minutes.

### Clamp conformation impacts commitment to termination principally through effects on elongation

We next set out to determine the effects of clamp conformation on intrinsic termination efficiency (TE) of ECs that elongate on templates containing one of three model intrinsic terminators: *t*_*hisL*_, λ*t*_R2_, and *t*_500_. (Figure 1A). By ligating one of three different linear DNA templates to the reconstituted ECs (Figure 3A), we could monitor the termination behavior at each terminator. We chose these terminators because they represent three classes of intrinsic terminators, as categorized by the composition and thus the strength of their U-tracts (Figure 1B): perfect (*t*_*hisL*_), near-perfect (λ*t*_R2_), and GC-containing (*t*_500_). Predictably and as reported previously, TEs varied significantly across the three terminators. Under reducing conditions, wild-type ECs exhibited a TE of 31 ± 3% at *t*_500_, 53 ± 6% at λ*t*_R2_, and 89 ± 2% at *t*_*hisL*_ (Figure 3C, 4A).

Despite the variability in U-tract composition and baseline TE, the general pattern of changes in TE after crosslink formation was consistent for λ*t*_R2_ and *t*_500_ (Figure 4A). Addition of diamide caused small reductions in the TE by the wild-type EC, despite its lack of a disulfide crosslink. Disulfide formation by D-LF greatly enhanced TE (36 ± 9% to 64 ± 6% at λ*t*_R2_; 23 ± 6% to 75 ± 2% at *t*_500_). Crosslinking of C-LF, on the other hand, depressed the TE at both terminators (38 ± 20% to 16 ± 3% at λ*t*_R2_; 19 ± 6% to 14 ± 2% at *t*_500_). C-259A responded to diamide treatment similarly to C-LF at all three terminators (*e.g.*, 41 ± 1% to 19 ± 4% at λ*t*_R2_) whereas the C-Δ3 EC appeared to behave more similarly to D-LF, exhibiting improved TE at both terminators (*e.g.*, 51 ± 4% to 63 ± 4% at λ*t*_R2_). These same trends were also observed at *t*_*hisL*_, but because the TE was too close to 100% (≥90%) the fold effects on TE were too small for robust quantitative analysis.

**Figure 4.**
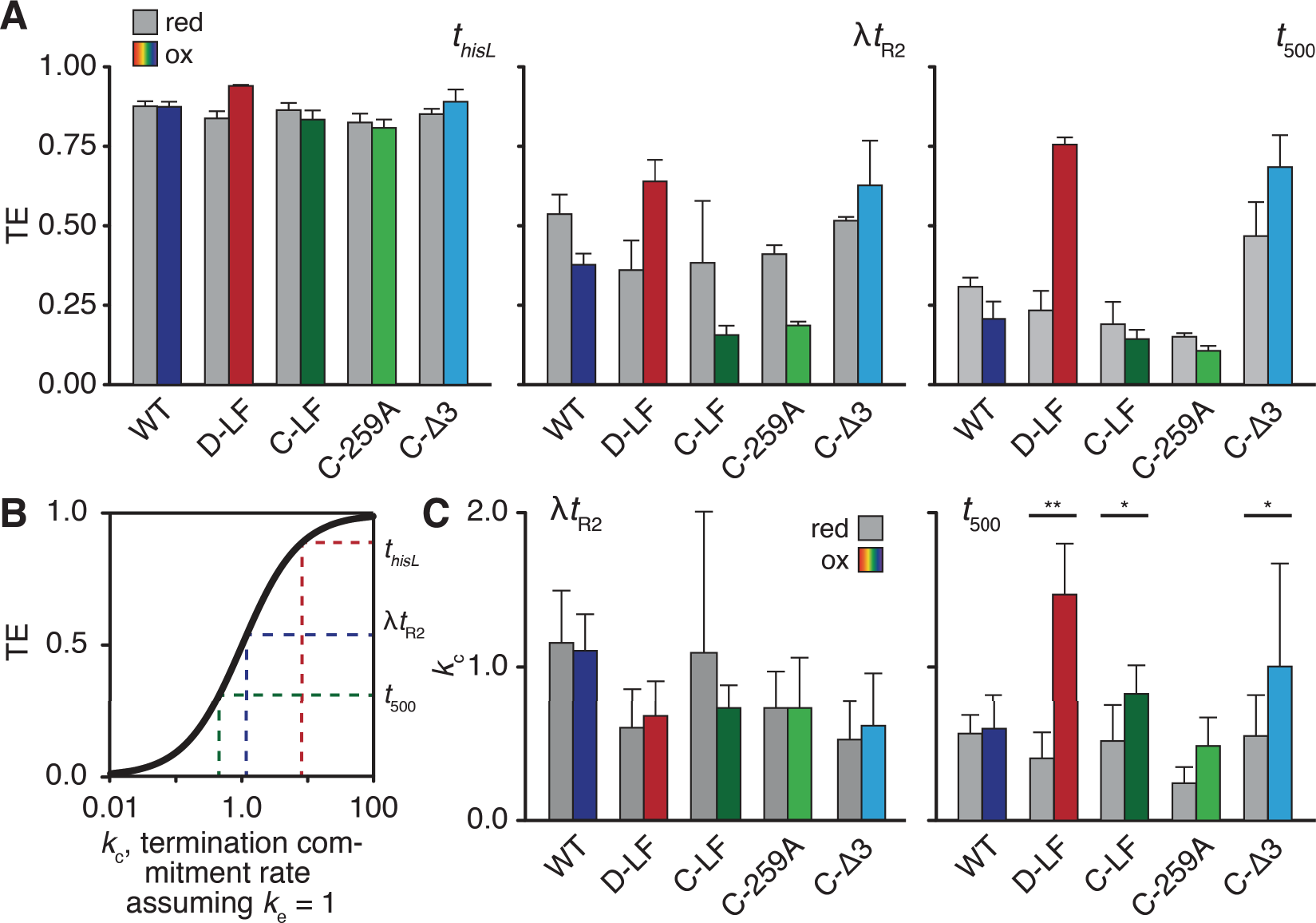
Evaluation of termination commitment rates. (**A**) Overall TE of Cys-pair RNAPs under reducing (grey bars) and oxidizing (colored bars) at three model terminators. Crosslinking efficiencies under these experimental conditions were: D-LF, 75±8%; C-LF, 73±5%; C-259A, 76±6%; C-A3, 81±11%. (**B**) TE is a sigmoidal function of the relative composite pseudo-first-order rates of elongation *k*_*e*_ and termination commitment *k*_*c*_, where TE = *k*_*e*_ / (*k*_*e*_ + *k*_*e*_). Shown are the effects on termination commitment rate, *k*_*c*_, where elongation rate *r*_*e*_, is set to 1. At low *k*_*c*_, small increases in rate result in significant jumps in TE whereas at high *k*_*c*_ even large increases will have small effects on TE. The high TE of *t*_*hisL*_ precludes accurate calculations of *k*_*c*_. (**C**) Conversion of TEs for λ*t*_R2_ and *t*_500_ into *k*_*c*_ where *k*_*c*_ = (TE × *k*_*e*_) / (1 - TE). As both TE and *k*_*e*_ were independent measurements with an associated s.d. (n = 3), error bars here represent the propagation of uncertainty (see Methods). *,*p* < 0.05; **,*p* < 0.01.

Comparison of the elongation rates of each enzyme with the TE revealed a general inverse correlation in which faster enzymes terminated less and slower enzymes terminated more. We therefore calculated relative commitment rates for each RNAP at λ*t*_R2_ and *t*_500_ using the simplifying assumption that EC inactivation, or termination commitment, is absolute so that TE is hyperbolically related to commitment rate (TE=*k*_c_/*k*_e_+*k*_c_, where *k*_e_ is the elongation rate and *k*_c_ is the commitment-to-termination rate; Figure 4B). At λ*t*_R2_, these analyses illustrated two key results (Figure 4C). First, each variant RNAP exhibited an intrinsic commitment rate (*k*_c_) distinct from that of the WT. Second, treatment with diamide and formation of a disulfide crosslink did not perturb this commitment rate significantly (*p* > 0.2 for all ECs), suggesting that restricting clamp movements does not impact the commitment process during termination.

However, the results were somewhat different at t_500_. Although treatment of the WT with diamide had little effect on commitment rate, formation of the D-LF, C-LF, C-Δ3, and C-259A crosslinks increased commitment rate, with the difference becoming statistically significant for D-LF (0.32 ± 0.13 to 1.17 ± 0.26; *p* > 0.01, Figure 4C). Taken together, these results suggest that lid-flap disulfides and clamp conformation can impact TE primarily by perturbing the transcription elongation rate, rather than affecting the termination pathway *per se*. However, for *t*_500_, which is thought to terminate *via* the forward translocation pathway due to the relative GC-richness of the U-tract [16], the lid-flap disulfides, notably D-LF, caused a modest additional effect on the termination commitment rate.

### RNA efficiently dissociates from an elongation complex at thisL regardless of clamp conformation

We next sought to explore how biasing clamp conformation with disulfide crosslinks and topologically closing the RNA exit channel or the main cleft affects the second step of the termination mechanism. Because measures of TE can only report on the commitment component of the termination mechanism, we modified the Cys-pair ligation scaffold assay to enable simultaneous study of RNA and RNAP release from bead-tethered DNA templates. As before, we reconstituted RNAP on an oligonucleotide scaffold upstream of a model terminator. We used *t*_*hisL*_ because its high TE allowed measurements of release; the lower TEs of λ*t*_R2_ and *t*_500_ precluded the reliable measurement of release behaviors. RNAP in ECs was radiolabeled by phosphorylation of a Protein Kinase A/Heart Muscle Kinase (PKA/HMK) tag sequence at the C-terminus of β′. Disulfide crosslinking was induced as before, after which all four NTPs were added to restart elongation and samples were removed at subsequent time points. At these time points, RNA or RNAP that was released from the paramagnetic bead-tethered DNA was collected after applying the reaction mixture to a magnetic rack for 15 s.

To close the RNA exit channel and inhibit clamp rotation or opening in the dissociation assay, C-LF was chosen because of its high crosslinking efficiency and its near-wild-type elongation rate under reducing conditions. To probe the effect of topologically closing the main cleft, we engineered a Cys-pair RNAP with cysteines in the β′ rudder and the β protrusion (T-RP; βA501C, β′S319C) that, when crosslinked, traps DNA within the cleft in addition to biasing the clamp towards the closed position (Figure S2). Treatment of T-RP with diamide induces formation of a disulfide within the transcription bubble, forming a steric block to release of DNA from the cleft. To promote release and make measurements feasible, we conducted assays in the presence of Cl^−^ (500 mM). Under these conditions, we achieved crosslinking efficiencies of 81 ± 4% for C-LF and 41 ± 5% for T-RP. By separating crosslinked and uncrosslinked, PKA-labeled β′ after completion of the reactions, we confirmed that the fraction of crosslinked ECs remained constant throughout the elongation and termination assay (Figure S5). The low crosslinking efficiency of T-RP may reflect the decreased ability of the Cys pair to fluctuate into reactive alignment or a suboptimal chemical environment for disulfide formation [36]. Under reducing conditions, both WT and C-LF ECs elongated more slowly (Figure 3B) but both RNAPs were able to elongate to *t*_*hisL*_ and terminate efficiently under these conditions within 20 s at 1 mM NTP (Figure S3). We therefore performed dissociation assays at 1 mM NTP.

By ^32^P-labeling both the nascent transcript and the β′ subunit of RNAP near its C-terminus, we could simultaneously monitor both RNA and RNAP dissociation from the bead-tethered DNA template because the RNA and β′ subunit electrophoretically separated to different regions of the gel (Figure S4). Under reducing conditions, RNA rapidly dissociated from the wild-type EC at efficiencies of greater than 85% by ~40 s (Figure S5A) and up to 95% after 2 min (Figure 5A). When the same assay was repeated under oxidizing conditions, only a slight decrease in the fraction transcript release was observed with the same rapid plateauing. We found that the bead release assay was subject to a relatively high experimental error that proved to be an unavoidable consequence of bead separation and supernatant recovery (Figure S5A–B). Although this error combined with the relatively rapid release on the time scale of the assay prevented accurate measurement of release rates, we could nonetheless draw basic conclusions whether release was fast or slow and about the efficiency of release using the plateau in release in the 120–480 s time window (Methods).

**Figure 5.**
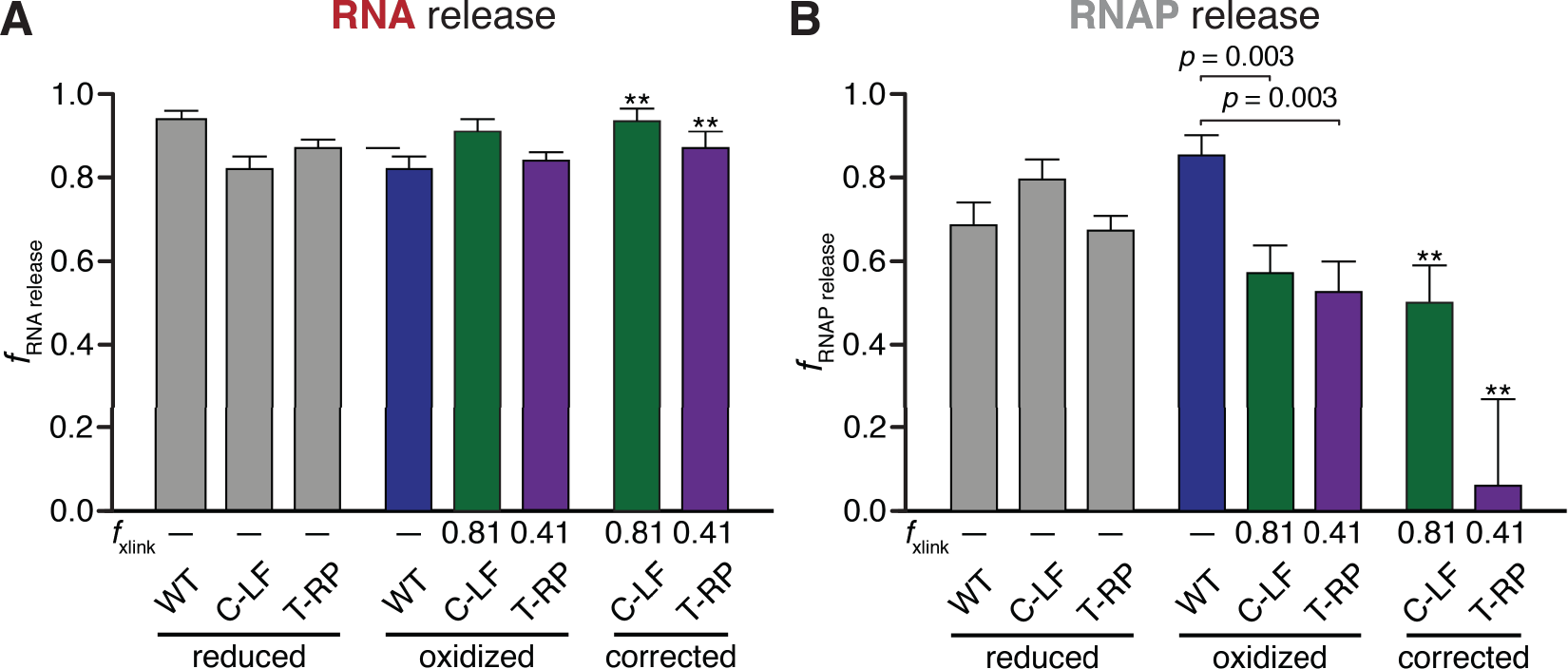
Evaluation of RNA and RNAP dissociation efficiencies. (**A**) Composite averages of RNA dissociation at *t*_*hisL*_ at 120, 240, and 480 s. Release was determined as the fraction of RNA released into the reaction supernatant relative to total RNA at the terminator site. Error represents the standard error of the mean (n=15-18). The independent effects of the disulfide crosslinks were determined by correcting RNA release values by C-LF and T-RP (see Methods). ** indicates propagation of error from standard deviations of crosslinking efficiency and s.e.m. measurements of RNA dissociation efficiencies. C-LF crosslinking efficiency, 81±4%; T-RP, 41±5%. (**B**) Composite averages of RNAP dissociation at *t*_*hisL*_ at 40, 50, and 60 s. RNAP dissociation was determined as the fraction of RNAP released into the reaction supernatant relative to total RNAP within the reaction mixture multiplied by the TE. Data was corrected as in subpanel A.

Using this approach, we found that treatment of WT ECs with diamide slightly decreased the efficiency of RNA release (from 94 ± 2% to 82 ± 3%) without large changes in the rate of release. For the Cys-pair crosslinks, we calculated the changes in efficiency after oxidation with diamide and also the corrected effect specific to the crosslinked fraction using the fraction crosslinked and the effect of diamide on WT ECs (rightmost set of bars, Figure 5A; see Methods). Both the C-LF and T-RP crosslinks minimally affected the fraction released RNA with or without correction for fraction crosslinked and had, at most, small effects on the rate of release (Fig. S5). Based on these results, we conclude that restricting the clamp in the closed conformation and covalently closing the RNA exit channel has at most a small delaying effect on RNA release at an intrinsic terminator. In other words, RNA can release out the exit channel even when clamp movement is restricted and the lid is crosslinked in place. These results are consistent with a model in which terminated RNA exits the RNA exit channel as single-stranded RNA after dissociation from the RNA–DNA hybrid without a need for large clamp movements.

### RNAP release from DNA is inhibited when clamp movement is blocked

Although clamp crosslinks had minimal effects on RNA release at the terminator, it remained difficult to envision how RNAP could dissociate from the DNA template without more dramatic movements based on available EC structures. Initiation is thought to require an open clamp to admit DNA into the main cleft [22, 37–39], making a requirement for clamp opening to release DNA logical.

Despite the challenges in quantification, the effects of clamp crosslinks on release of RNAP from DNA were obvious (Figure 5B). Examination of RNAP release prior to 100 s reveals that treatment of the wild-type EC with diamide actually *increased* the efficiency of release (69 ± 6% to 86 ± 5%; Figure 5B). This increase in RNAP release is consistent with oxidized RNAP binding the NA scaffold somewhat more weakly, which could also explain its faster elongation rate (see Figure 3B). In contrast, the C-LF and T-RP crosslinks both inhibited RNAP release from DNA. This effect is most obvious from the decrease in the total amount of RNAP released, which was highly significant relative to wild-type even before correcting for the previously noted effects of oxidation on the EC (*p* = 0.003 for both C-LF and T-RP; Figure 5B). After correcting for the fraction crosslinked (see Methods), we found that C-LF and T-RP release efficiencies decreased to 50 ± 9% and 6 ± 21%, respectively. These results suggest that crosslinking the clamp to either the flap or the protrusion strongly inhibits release of RNAP from DNA even though the RNA transcript can still release efficiently. This inhibition may reflect either the narrow width of the RNAP cleft when the clamp is closed, which would sterically block DNA release, or contacts of the clamp and switch regions to DNA that inhibit release unless the clamp opens. Regardless, RNA release without DNA release appears to be possible at an intrinsic terminator. This result has important implications for both the mechanism of termination and the fate of RNAP after termination (see Discussion).

### Inhibiting RNAP conformational flexibility with microcin J25 impairs both RNA and RNAP release

Kinetic analyses led to the development of a multistate-multipath model of termination commitment in which restricting movements of the polymorphous TL motif slows the rate of termination by reducing the probability that ECs positioned at a terminator fluctuate by thermal motion into a state committed to termination [14]. However, experiments to date only probed possible effects of TL restriction on the commitment step and not the release step of termination. Our current clamp crosslinking results suggest that large motions of the clamp are not required for either irreversible commitment to termination or RNA release. To ask if TL-mediated fluctuations other than large clamp movements are required for release, we tested the addition of microcin J25 (MccJ25). MccJ25 is a 21-residue antimicrobial lariat peptide [40] that binds and blocks the secondary channel, an element of RNAP thought to act as an “access tunnel” for incoming NTPs to enter the enzyme’s active site, as a regulator binding site, and as a path for RNA during backtracking (Figure 6A). Based on the location of resistance mutations, MccJ25 is also thought to contact the TL, blocking catalysis by inhibiting TL motion [41–43]. We prepared MccJ25 and used an abortive initiation assay (Figure S6) to verify its activity. Under our experimental conditions, we determined that this preparation of MccJ25 inhibited abortive initiation at P*lac*UV5 with an IC_50_ of 0.63 ± 0.03 μM (Figure S6A), close to the published *K*_i_ of 1.4 µM [41].

**Figure 6.**
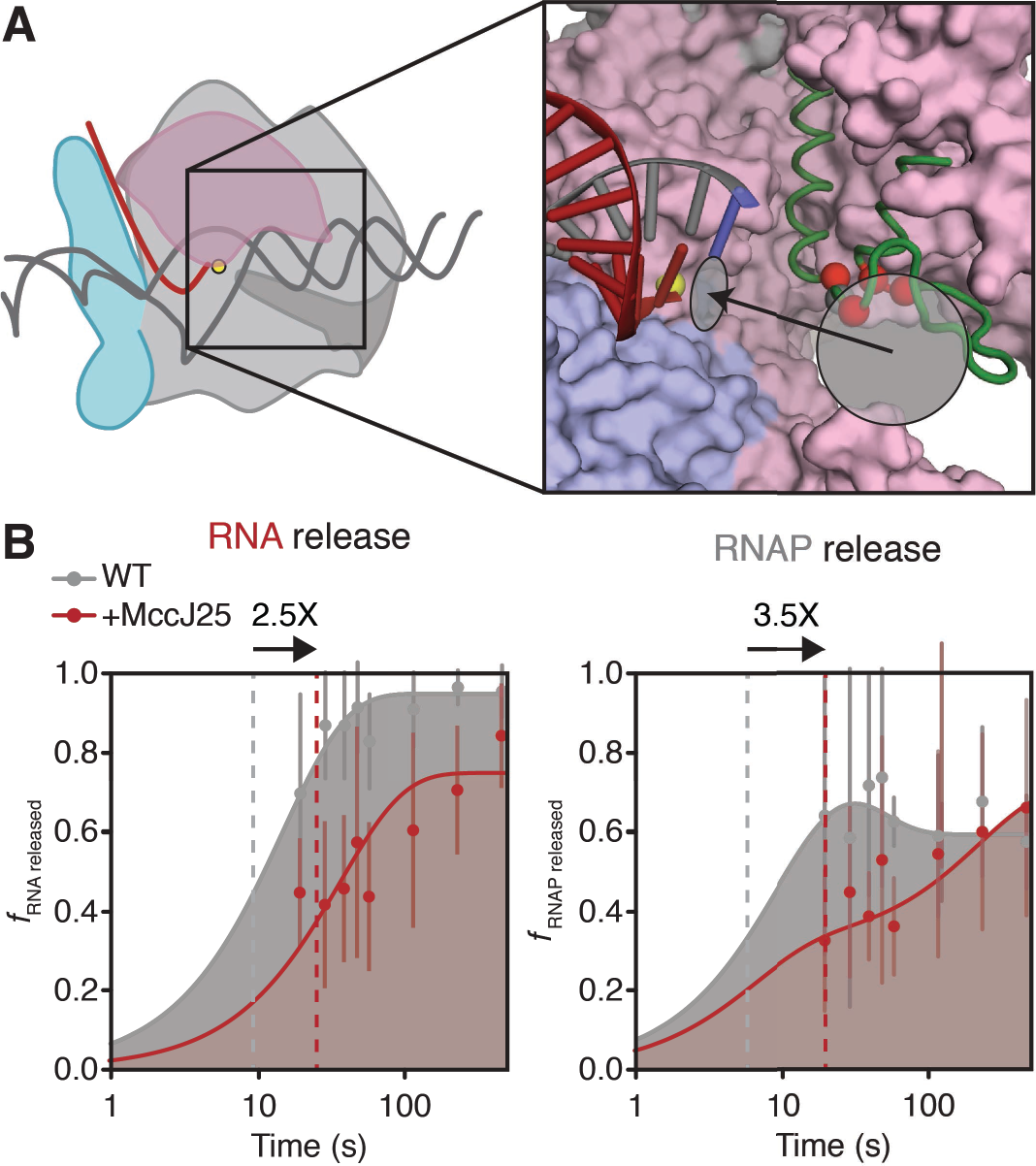
Evaluation of RNA and RNAP dissociation rates in the presence of MccJ25. (**A**) MccJ25 bind the TL and the RNAP secondary channel. Red spheres indicate the Ca atoms of TL residues that confer MccJ25 resistance in mutational screens [41]. The grey sphere indicates the Cα atom of β′T931. The light grey circle indicates the location of the RNAP secondary channel and the black arrow indicates the path of NTP into the active site, light grey ellipse. The RNA-DNA hybrid is shown; other nucleic-acid components are omitted for clarity (red, RNA; grey, DNA; blue, template DNA base positioned in the RNAP active site). Core RNAP components are derived from PDB ID: **<u>6AAF</u>** [27]; green, unfolded TL, PDB ID: **<u>2Q5J</u>**. (**B**) Semi-log plots of RNA and RNAP release rates at *t*_*hisL*_ When relevant, MccJ25 is added 5 s after addition of chase NTPs. RNA dissociation was determined as the fraction of RNA released into the reaction supernatant relative to total RNA at the terminator site; RNAP dissociation was determined as the fraction of RNAP released into the reaction supernatant relative to the amount of total RNAP within the reaction mixture multiplied by the TE. Grey curves are the same as those in Figures S5A-B; red curves represent RNA and RNAP release from WT RNAP in the presence of 100 μM MccJ25. Dashed lines represent the time at which half of the maximal released RNA was achieved over the time course. Arrows on top of plots represent the fold-change in the time it takes for half-maximal release to occur upon treatment with MccJ25. Error bars represent the standard deviation at n = 3.

We next used an established disulfide reporter assay that detects different TL conformations [36, 44] to determine whether MccJ25 restricts TL conformations (Figure S7A). Because genetically identified binding determinants for MccJ25 overlap the TL, including the location of strong resistance mutations at TL residue T931, we anticipated that MccJ25 would limit TL movement and thus reduce crosslinking of the disulfide reporters. As expected, incubation of RNAP with MccJ25 severely reduced the crosslinking efficiency of Cys-pairs that detect the fully-folded trigger helices and a partially-folded conformation (Figure S7B). MccJ25 effects on two Cys-pairs that report the unfolded conformation, however, were smaller, suggesting that MccJ25 positions the TL closer to unfolded conformations that enable these crosslinks but may also reduce TL flexibility. The reduction in crosslinking by all four reporters of TL conformation confirms that MccJ25 strongly constrains TL conformational fluctuations.

Having established that MccJ25 blocks TL motion, we next asked if inhibiting TL motions would affect the rate of RNA or RNAP release from DNA at *t*_*hisL*_. We evaluated the effects of MccJ25 on dissociation at *t*_*hisL*_ by allowing rapid elongation of A26 ECs to the terminator in the high NTP conditions used for the release assays and then adding MccJ25 after 5 s. Addition of MccJ25 at 5 s allowed enough time for most ECs to reach the terminator prior to inhibition of elongation by MccJ25, as indicated by a negligible effect on the RNA products present in the reaction (Figure S6B), but was still prior to the release of most RNA and RNAP at the terminator (Figure S5). This approach enabled us to assay for the effect of MccJ25 on release in the 20–480 s time window (Figure 6). Based on the shift in the time of half-maximal release, MccJ25 slowed the rate of release of RNAP from template DNA by a factor of 3.5 Strikingly, addition of MccJ25 also slowed the release of the nascent transcript by a factor of 2.5 (Figure 6B). We conclude that TL motions increase the rate of RNA and RNAP release at an intrinsic terminator in addition to their roles in steps leading to commitment.

## DISCUSSION

Our study provides new understanding of the role of RNAP conformational changes in the mechanism of intrinsic termination. By developing a bead-tethered, ligated-scaffold assay, we could disentangle the steps of transcription elongation, commitment, and dissociation that are conflated in the traditional measurement of TE and thereby gain several key insights. First, although clamp conformation affects TE, these effects are due primarily to changes in the elongation rate rather than changes in the commitment rate except at GC-containing imperfect terminators (*e.g*., *t*_500_) that release RNA by forward EC translocation rather than hybrid shearing. Second, we found that clamp movements are more important for dissociation of RNAP from DNA than for dissociation of RNA from the EC, which is the defining step in termination. Finally, we found that restricting movements of the TL markedly slowed the rates of both RNA and RNAP release, consistent with a role for TL-mediated conformational fluctuations in the termination process [14].

### Clamp conformation has stronger effects on elongation than on termination

Structural and biochemical analyses of ECs suggest roles of the clamp in promoting EC stability *via* numerous non-specific interactions with the NA scaffold [26], in initiation by opening and closing to admit DNA to the cleft [22, 39], and in hairpin-stabilized pausing by aiding the swiveling that inhibits TL folding and thus catalysis [30, 33]. Here we find that clamp conformation affects the efficiency of intrinsic termination, but primarily through effects on the competing rate of elongation rather than in the termination pathway *per se*. A disulfide bond that biased the clamp toward the displaced (swiveled) conformation observed in paused ECs (D-LF), or a small deletion in the lid (C-Δ3) slowed elongation whereas stabilizing the clamp in the closed conformation (C-LF) accelerated elongation. These effects on elongation, all of which caused corresponding changes in TE, are likely caused primarily by changes in pausing, which were evident in transcription elongation patterns (Figure S1). Although we cannot exclude effects on on-pathway elongation, it is notable that the D-LF and C-LF disulfides that affect intrinsic termination only indirectly have large and opposite effects on a canonical hairpin-stabilized pause [31], and that pausing is known to occur at intrinsic terminators [10, 14].

The key point is that clamp conformation affects pausing/elongation much more than it affects the pathway of RNA release at an intrinsic terminator, which has important implications. Bacterial intrinsic terminators have apparently evolved to be robust relative to EC conformation. Whereas pausing and the overall rate of elongation may be readily modulated by clamp-binding elongation factors (*e.g.*, RfaH, NusG, Spt4/5, hepatitis δ antigen; [45–47]), the insensitivity of canonical bacterial terminators to clamp conformation may enable intrinsic terminators to halt transcription irrespective of EC composition. A case in point may be the *E. coli* ribosomal RNA operons (*rrn*), which are transcribed by antitermination-modified ECs [48, 49] and end with intrinsic terminators that contain G-C bp in their U tracts and thus appear to be in the forward-translocating class. Antitermination proteins may limit clamp and lid mobility similarly to the disulfide crosslinks making termination efficient at the forward-translocation-dependent *rrn* terminators (see discussion of a possible mechanistic basis for this effect below).

The lack of inhibition of RNA release by lid–flap crosslinking also provides further reason to doubt the hairpin-TL contact model on intrinsic termination [19]. This model, which also has been questioned on other grounds [14–17, 50], requires the terminator hairpin disrupt RNAP interactions by sweeping through the cleft, starting by breaking the lid-flap interaction. Since RNA release is largely unaffected when the lid is crosslinked to the flap, this model seems problematic.

Deletion of three residues from the tip of the lid loop (C-Δ3) slowed elongation/increased pausing even under reducing conditions that disfavored a disulfide that constrained the clamp. A close inspection of recent structures of the *Escherichia coli* EC suggest that this effect may be due to remodeling of the upstream fork junction, where the RNA-DNA hybrid is separated, single-stranded RNA is threaded into the RNA exit channel, and template DNA reanneals with the nontemplate strand [27]. The C-259A variant RNAP demonstrated that a single substitution in this region did not significantly perturb elongation, but it is likely that deletion or mutation of all three of these conserved residues would have additive or synergistic effects on the architecture of the fork junction. The deleted residues may ordinarily guide strand exchange and thus translocation at the fork junction. Thus, the C-Δ3 deletion may slow elongation or stimulate pausing by inhibiting translocation.

We also provide evidence that the general redox state of the EC impacts its elongation rate, irrespective of the presence of a disulfide crosslink. Treatment of the wild-type EC with diamide increased its overall elongation rate nearly two-fold. RNAP contains 28 cysteine residues, eight of which play a role in binding Zn^2+^ ions, and one of which is in close proximity to the active site (*Eco* β′C454). It is possible that chemical modification of any of these cysteines could impact the elongation rate of the EC by altering RNAP conformation in a way that stimulates catalysis (*e.g.*, promoting TL folding) or by indirectly affecting NA interactions. These results highlight the need for in-depth investigation into the role of redox state on RNAP.

### Upstream fork-junction mobility may weaken terminators that require forward translocation for commitment

EC inactivation/commitment at terminators for which hybrid shearing is a viable route was unaffected by biasing clamp conformation with disulfide crosslinks (*t*_*hisL*_ and λ*t*_R2_; [16]). However, this process was more affected by crosslinks for a terminator that uses forward translocation without nucleotide addition as the path to EC inactivation/commitment (*t*_500_; Figure 4B). This result suggests that hybrid shearing can occur without major changes in clamp conformation and that for hybrid-shearing terminators, any role of a hairpin-stabilized pause intermediate must be simply to increase the time window for shearing to occur and not in the inactivation process itself (these crosslinks are known to affect hairpin-stabilized pausing [30]). That a forward-translocation terminator exhibits different responses to the disulfide crosslinks further underscores the existence of an alternative route to termination at terminators with GC-containing U-tracts [18].

The effect of disulfide crosslinks at terminators with GC-containing U-tracts appears more related to restricting mobility of the lid than a role of any particular clamp conformation. We draw this conclusion because crosslinks that bias the clamp toward either displaced (D-LF) or closed (C-LF) conformations both increase the rate of commitment at *t*_500_ (Figure 4C), indicating that no particular clamp conformation favors or disfavors forward translocation and instead that restricting lid mobility in any state increases termination. Additionally, the non-crosslinked form of the lid Cys insertion combined with R259A decreased termination at *t*_500_, further indicating that the lid may influence this class of terminators.

Formation of a crosslink by the C-Δ3 RNAP and disruption of correct upstream fork junction architecture also increased the commitment rate at *t*_500_. C-Δ3 was constructed by removing β′L255, D256, and R259, all of which are conserved across diverse species and, as discussed above, may be involved in translocation. L255 interacts with the −9 base pair of the hybrid, D256 forms a salt bridge with βR841 in the flap to seal the RNA exit channel, and R259 forms a base stack with the −10 base of the template strand [27]. Each of these interactions plays a role in forming the proper geometry of the upstream fork junction and have been observed to shift upstream with the NA scaffold in a translocation intermediate [30]. Deletion of these residues could have additive or synergetic effects with restricting lid movements on hypertranslocation.

In support of this hypothesis, the lid appears to have pronounced effects on forward translocation of RNAP adjacent to a terminator hairpin [51]. Based on these observations, we speculate that restricting lid mobility with disulfide crosslinks may destabilize the upstream fork-junction because lid mobility stabilizes the hybrid as it shifts translocation register. For terminators like *t*_500_ that require forward translocation unlinked to nucleotide addition to commit to termination, this hybrid destabilization may aid hybrid separation during forward translocation and increase the overall rate of commitment.

### Clamp opening appears to be required for RNAP release from DNA at intrinsic terminators

It is structurally plausible for an EC to become catalytically inactivated without dramatic movements of the clamp. However, it is less easy to understand how RNAP could dissociate from template DNA without clamp movements. Reannealing of the transcription bubble would generate an unmelted DNA duplex that would be pulled out of the active site and be too large and straight to fit within the main cleft of RNAP in either the closed or swiveled clamp conformation. Thus, current structural understanding of the EC requires large movements of the clamp along the axis perpendicular to the hybrid to allow the dissociation of RNAP from the DNA [22]. However, the same structural data suggest that such large-scale clamp motions should not be necessary for release of the nascent transcript out of the exit channel once the hybrid is shortened to an unstable length.

Our studies of the dissociation step in termination support these views. Formation of a crosslink that stabilized a closed clamp had at most minor effects upon the rate or efficiency of RNA release at an efficient terminator (Figures 5A and S5A). However, both the efficiency and, qualitatively, the rate at which RNAP dissociated from the DNA at the terminator were reduced under conditions that disfavor clamp displacement (Figures 5B and S5B), suggesting a need for orthogonal opening of the clamp during dissociation. These effects were most dramatic when a rudder-protrusion crosslink was formed within the transcription bubble to induce a steric block to complete reannealing of duplex DNA (Figure 5B). Taken together, these results indicate that whereas RNA can easily shear out of the RNA exit channel at *t*_*hisL*_ even when clamp movement is constrained by a disulfide, RNAP release from template DNA depends both upon the clamp’s ability to move orthogonally from the main cleft (*i.e*., to “open”) and upon the template strand’s ability to reanneal with the non-template strand in order to induce collapse of the transcription bubble [52].

A further implication of our finding that RNA release at a terminator can occur without RNAP release from DNA is that RNAP may remain bound to DNA after termination in some cases. If an unreleased RNAP were still associated with a σ initiation factor, as has been observed to occur in some situations [53–55], then it may be able to enter a promoter search mode directly without release from the DNA.

### TL movement is required for release of both RNA and DNA at an intrinsic terminator

Previous studies of the role of the TL in intrinsic termination mechanism led to a model in which termination was promoted not by a TL contact to the terminator hairpin but by conformational fluctuations of the TL that allowed an EC located at a terminator to sample multiple conformations [14]. This multistate-multipath mechanism was proposed as an extension to the classical thermodynamic model of termination as a competition between elongation *vs.* termination [5]. Our findings that restricting the movements of the TL with the antimicrobial peptide MccJ25 slows the dissociation rates from template DNA are consistent with the multistate-multipath model, although they do not distinguish effects on commitment *vs.* dissociation. Because we added MccJ25 to ECs already located at a terminator but not yet dissociated, because MccJ25 would inhibit subsequent elongation, and because our readout was release of RNA or RNAP, it is possible that the effects of restricting TL movements with MccJ25 could be manifest at either the commitment or dissociation steps in the mechanism. Regardless, these effects are striking and suggest that restricting TL movement stabilizes ECs located at a terminator, disfavoring termination. During termination commitment, the 3′-end of the RNA transcript should be displaced from the active site of RNAP by at least 2–3 base pairs [17], positioning the last base several Ångstroms from the TL in either its folded or unfolded conformation. Possibly, restricting TL movements slows this displacement, as the folded form of the TL is thought to contact or at least stabilize RNA in the pretranslocated register [56]. Alternatively, stabilizing the TL in an open conformation may allow the initially paused RNA at a terminator to backtrack into the secondary channel; such an effect would slow both RNA and RNAP release.

In either scenario, limiting TL movement may reduce thermal motions of the complex set of interconnected modules in RNAP whose normal fluctuations aid release of RNA and DNA from RNAP (the core idea in the multistate-multipath model). If conformational restrictions alone account for slowing termination, then these limitations apparently are much greater when MccJ25 is bound than when the clamp is restricted by a disulfide crosslink since the former but not the latter slows RNA release.

### Conclusion

Our findings support the idea that the release step of termination mirrors initiation in requiring clamp movement to allow DNA escape from the main cleft of RNAP just as it is required for DNA entry during initiation. The fact that RNA can release and transcription can terminate when RNAP clamp opening is inhibited may help explain how intrinsic terminators can function when elongation or antitermination factors stabilize the EC conformation. The possibility that RNAP could remain bound and enter a nonspecific sliding mode on DNA without fully dissociating at a terminator merits further study.

## MATERIALS AND METHODS / EXPERIMENTAL PROCEDURES

### Materials

All DNA and RNA oligonucleotides were obtained from IDT (Table 1). Oligonucleotides used for reconstitution of elongation complexes were gel purified by denaturing polyacrylamide gel electrophoresis (PAGE). HPLC-purified NTPs were from Promega Corp. [α-^32^P]-GTP and [γ-^32^P]-ATP were from Perkin-Elmer. T4 Sty I-HF and 2,000,000 U T4 DNA Ligase/ml were from New England Biolabs.

**Table 1.**
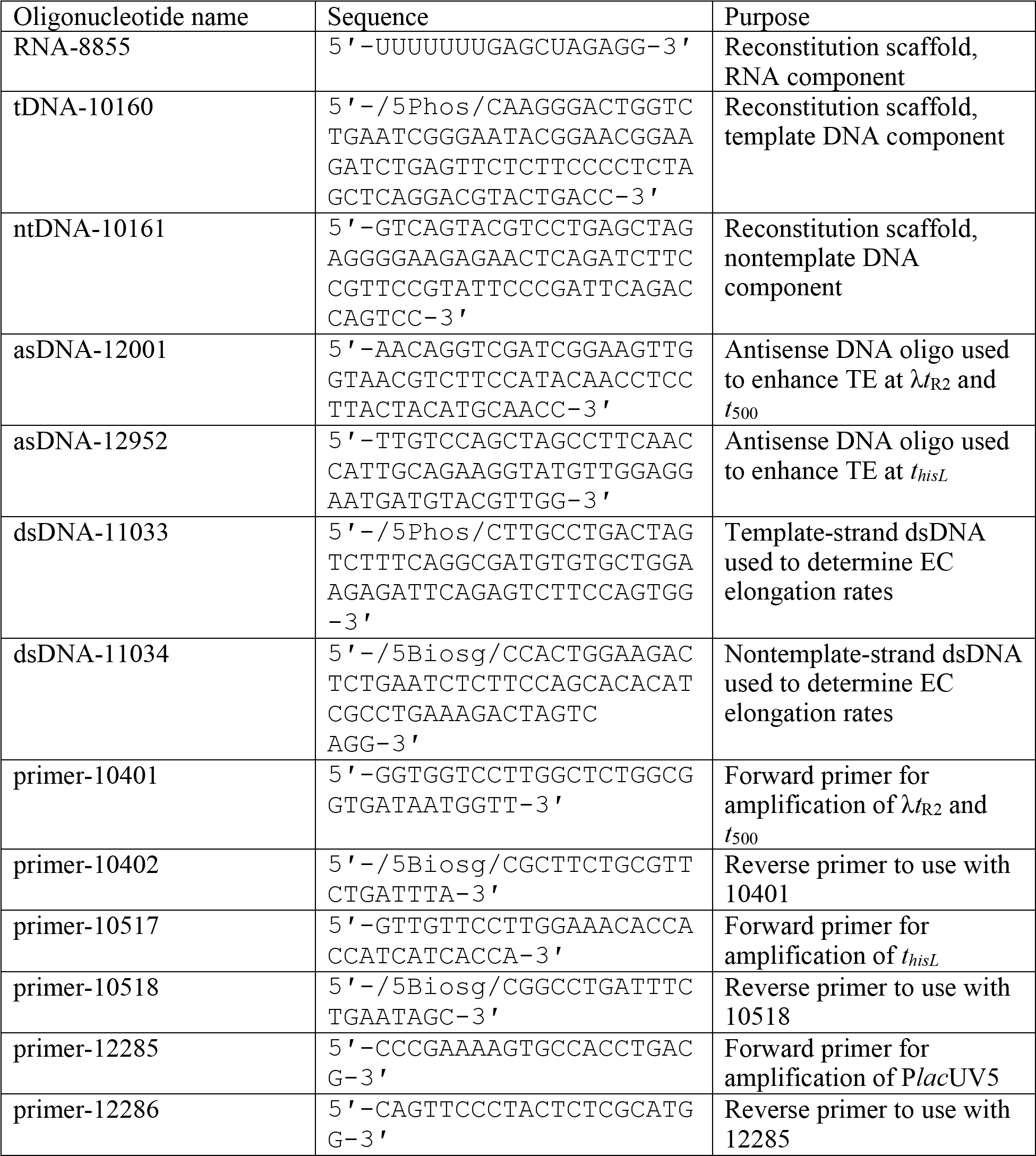
Oligos

### Protein and peptide purifications

Wild-type RNAP, Cys-pair RNAPs, and wild-type σ^70^ were overexpressed in T7 Express competent *E. coli* (New England Biolabs) from plasmids (Table 2). The parent RNAP overexpression plasmid pRM843 is a T7 RNAP-driven overexpression plasmid that encodes *E. coli rpo*A, *rpo*Z, *rpo*B and *rpo*C, yielding core RNAP. The N-terminus of β contains a His_10_ tag followed by a PreScission protease cleavage recognition signal (LEVLFQGP). The C-terminus of β′ contains a *Strep*-tag^Ⓡ^ preceded by a Protein Kinase A (PKA) recognition signal (GRTGRRNSI). Competent *E. coli* cells were transformed and RNAP was overexpressed and purified as previously described [31, 57] except that the purified product was dialyzed into a high [DTT] RNAP storage buffer (10 mM Tris-Cl, pH 7.9, 20% [v/v] glycerol, 100 mM EDTA, 10 mM DTT, 100 mM NaCl, 1 mM MgCl_2_, 20 μM ZnCl) to ensure the integrity of the cysteine sulfhydryl groups. Aliquots were stored at −80 °C. RNA polymerase holoenzymes used in MccJ25 abortive initiation control assays were formed by incubating 2 μM purified core enzyme with 4 μM σ^70^ in RNAP storage buffer, incubated at 30 °C for 30 minutes, and stored at −20 °C before use.

**Table 2.** Plasmids

| Plasmid name | Source | Purpose |
| --- | --- | --- |
| pRM843 | [31] | Overexpression of wild-type RNAP (*rpoA*, *rpoZ* His<sub>10</sub>:PPX:*rpoB*, *rpoC*:PKA:*strep*) |
| pRM890 | [31] | pRM843 derivative. Overexpression of D-LF RNAP (βT843C $\beta'$258iC) |
| pRM918 | [31] | pRM843 derivative. Overexpression of C-LF RNAP (βP1044C $\beta'$258iC) |
| pRM1100 | This paper | pRM843 derivative. Overexpression of C-Δ3 RNAP (βP1044C $\beta'$ Δ255–256,258iC,Δ259) |
| pRM1177 | This paper | pRM843 derivative. Overexpression of C-259A RNAP (βP1044C $\beta'$258iC R259A) |
| pRM1001 | This paper | pRM843 derivative. Overexpression of T-RP RNAP (βA501C $\beta'$S319C) |
| pDJ115 | [36] | Overexpression of P937-687 ($\beta'$ I937C P1135C:PKA:His<sub>10</sub>) |
| pDJ116 | [36] | Overexpression of P937-687 ($\beta'$ I937C T1135C:PKA:His<sub>10</sub>) |
| pDJ124 | [36] | Overexpression of F937-746 ($\beta'$ Q736C I937C:PKA:His<sub>10</sub>) |
| pDJ129 | [36] | Overexpression of P937-687 ($\beta'$ G687C I937C:PKA:His<sub>10</sub>) |
| pET-28-$\sigma^{70}$ | Gift from R.B. | Overexpression of $\sigma^{70}$ (His<sub>6</sub>:*rpoD*) |
| pJP73 | Gift from A.J.L. | Overexpression of MccJ25 |
| pIA171 | [64] | Plasmid template for PCR amplification of *thisL* |
| pRM1065 | This paper | Plasmid template for PCR amplification of $\lambda t_{r2}$ |
| pRM1066 | This paper | Plasmid template for PCR amplification of $t_{500}$ |

His-tagged wild-type *E. coli* σ^70^ was overexpressed in TE Express from the pET-28a-σ^70^ overexpression plasmid (a gift from Richard Burgess, University of Wisconsin–Madison, WI) by growing to an OD_600_ of ~0.5 and induced by addition of IPTG to 0.4 mM. The temperature was decreased to 16 °C and cells left shaking at 200 rpm overnight. Cells were harvested by centrifugation at 3000 ×g for 15 min at 4 °C followed by sonication in lysis buffer (50 mM Tris-Cl, pH 7.9, 5 mM imidazole, 5% [v/v] glycerol, 233 mM NaCl, 2 mM EDTA, 10 mM β-mercaptoethanol) plus a protease inhibitor cocktail (final concentrations 100 μg phenylmethylsulfonyl fluoride/ml, 0.624 mg benzamide/ml, 10 μg chymostatin/ml, 10 μg leupeptin/ml, 2 μg pepstatin/ml, 20 μg aprotonin/ml, 20 μg antipain/ml). The lysate was centrifuged at 11000 ×g for 15 min at 4 °C and the σ^70^-containing supernatant was filtered and applied to a 5-ml HisTrap HP column (GE Healthcare). Protein was eluted over a gradient of 5– 500 mM imidazole in lysis buffer and fractions were analyzed on 20% SDS-PAGE. σ^70^-containing fractions were pooled and run through a HiLoad 16/60 Superdex 200 prep grade (GE Healthcare) size exclusion column. Purified σ^70^ was dialyzed overnight into σ^70^ storage buffer (10 mM Tris-Cl, pH 8.0, 30% [v/v] glycerol, 0.1 mM EDTA, 100 mM NaCl, 20 μM ZnCl_2_, 1 mM MgCl_2_, 0.1 mM DTT) and aliquots were stored at −80 °C.

MccJ25 was purified from the pJP73 overexpression plasmid (a gift from A. James Link, Princeton University, NJ) by a protocol adapted from [58]. Briefly, DH10B cells containing the pJP73 plasmid were grown up in LB overnight at 37 °C at 250 rpm. Cultures were diluted to a starting OD_600_ of ~0.02 in M9 salts supplemented with thiamine, glucose and all 20 amino acids, and grown at 37 °C at 250 rpm to a final OD_600_ of 0.5. The *mcj*ABCD operon was then induced with 1 mM IPTG and grown overnight for ~20 hrs at 37 °C at at 250 rpm. Cells were pelleted and MccJ25-containing supernatant collected. MccJ25 was purified from the supernatant on BondElut C8 cartridges (Varian, now acquired by Agilent Technologies) following manufacturer instructions. The elutions were dried using a rotary evaporator, followed by a Speedvac, and then dissolved in water. To check purity, products were resolved on a homogenous 20% denaturing PhastGel (GE Healthcare), followed by Coomassie brilliant blue R-250 staining.

### Producing linear DNA fragments for transcription

Linear DNA templates were generated from plasmids *via* PCR (Table 1 & 2) using OneTaq^Ⓡ^ DNA polymerase (New England Biolabs) and purified *via* spermine precipitation [59]. Briefly, PCR products were incubated with 10 mM spermine at 4 °C for 15 minutes followed by high-speed centrifugation and removal of the supernatant. Samples were incubated with spermine exchange buffer [75% (v/v) ethanol, 300 mM sodium acetate, 10 mM Mg(OAc)_2_] at 4 °C for 1 hr followed by another high-speed centrifugation for 5 min. The supernatant was removed and the pellet was washed with 70% (v/v) ethanol and centrifuged as before. Ethanol was evaporated off *via* speed-vac and the pellet was dissolved in TE. Sample concentration was measured using a NanoDrop Spectrophotometer (Thermo Fisher Scientific). DNA fragments were processed with StyI-HF restriction endonuclease following the manufacturer’s protocol and cleaned through a second round of spermine precipitations.

### Transcription on fully complementary oligonucleotide scaffolds

ECs were reconstituted step-wise. First, excess template strand (10 μM) was annealed to 5 μM RNA in reconstitution buffer [10 mM Tris-OAc, pH 7.9, 40 mM KOAc, 5 mM Mg(OAc)_2_] by heating to 75 °C for 2 min, rapidly cooling to 45 °C, and then cooling to room temperature in 2 °C/2-min steps. This initial scaffold was diluted to 500 nM by addition of elongation buffer (EB; 20 mM Tris-OAc, pH. 8.0, 40 mM KOAc, 5 mM Mg(OAc)_2_, 0.1 mM EDTA, 1 mM DTT) followed by addition of 2.5 μM RNAP and incubation at 37 °C for 15 min. Non-template strand was then added to 1.5 μM and the sample was again incubated at 37 °C for 15 min, yielding an EC formed with the molar ratio 1 RNA:2 tDNA:3 ntDNA:5 RNAP. PCR fragments were pre-bound to paramagnetic beads (Thermo Fisher Scientific) through a 5′ biotin moiety on the template strand. Next, 150 nM bead-bound fragments were mixed with 50 nM ECs at a 3:1 ratio, and ligated together with 2,000 U T4 DNA ligase and 1 mM ATP in EB at 16 °C for 2 hr. Non-ligated ECs and excess RNAP were removed from the mixtures through two EB washes on a magnetic rack. RNAs were labeled by addition of 0.2 mCi [α-^32^P]-GTP and heparin was added as a competitor to 50 μg/ml. Disulfide crosslinking of the Cys-pairs was initiated by adding diamide to 15 mM and incubating ECs at 37 °C for 15 min. To determine crosslinking efficiencies, aliquots were removed, quenched with 15 mM iodoacetamide, and separated through SDS-PAGE on 8% Bolt gels in 1X MOPS Bolt running buffer (Thermo Fisher Scientific,), stained with Krypton Flourescent Protein Stain (Thermo Fisher Scientific) according to the manufacturer’s protocol, imaged with a 520-nm excitation filter and 580-nm emission filter, and quantitated *via* ImageQuant. The remaining sample was diluted in EB ± oxidant to a working concentration of 5 nM, and a DNA oligonucleotide antisense to upstream sequences was added to 1 mM to prevent formation of secondary structures that could block T_hp_ formation. Transcription was initiated by saturation with 150 μM ATP, UTP, and CTP and 30 μM GTP. During endpoint assays, reactions were stopped at 10 min by addition of an equal volume of 2X urea stop buffer (90 mM Tris-borate, pH 8.3, 8 M urea, 50 mM EDTA, 0.02% [w/v] xylene cyanol, 0.02% [w/v] bromophenol blue). In timepoint assays, reactions were likewise stopped at 15, 30, 60, 120, 240, 480, 960, and 1920 s. Samples were separated by electrophoresis in a denaturing 8% (v/v) polyacrylamide gel and quantitated after imaging with a PhosphorImager.

### Dissociation assays

Except where noted, release assays were performed using complexes formed as described above. To promote dissociation, RNA and template-strand DNA were annealed in Cl^−^ reconstitution buffer (10 mM Tris-Cl, pH 7.9, 40 mM KCl, 5 mM MgCl_2_) and all subsequent steps were performed in Cl− elongation buffer (20 mM Tris-Cl, pH. 8.0, 40 mM KCl, 5 mM MgCl_2_, 0.1 mM EDTA, 1 mM DTT). The Protein Kinase A (PKA) tag on the β′ CTD was radiolabeled by addition of 7.5 U PKA and 0.5 mCi [γ-^32^P]-ATP/μl at the same time [α-^32^P]-GTP was added for incorporation labeling of the RNA. This step was performed after reconstitution to block access of PKA to the rifampicin-binding pocket, which can serve as a second labeling site on RNAP [60]. To reduce the possibility of aggregation of released RNAPs, 1 μM double-stranded DNA was used as a competitor instead of heparin. Antisense DNAs were not added in these assays. Transcription was restarted by addition of 500 mM NaCl and all four NTPs to 1 mM. During MccJ25 studies, MccJ25 was added to 100 μM 5 s after NTPs. Reactions were stopped with an equal volume of 2X urea/SDS stop buffer (90 mM Tris-borate, pH 8.3, 8 M urea, 4% [w/v] SDS, 50 mM EDTA, 0.02% [w/v] xylene cyanol, 0.02% [w/v] bromophenol blue), separated by denaturing 8% SDS Bolt gels, and imaged and quantitated using a PhosphorImager. Release efficiencies at individual timepoints were calculated as the signal of RNA or RNAP released into the supernatant divided by the total signal of released and bead-bound RNA or RNAP (Figure S4). RNAP release values were normalized to the overall termination efficiency. Next, composite averages of samples taken at 40, 50, and 60 seconds (for RNAP) and 120, 240, and 480 s (for RNA) were calculated. We used the peak release at 40–60 s for RNAP because aggregation or non-specific effects reduced RNAP in the supernatant at later time points. As a result, it is possible that some of the apparent effect on fraction RNAP released could reflect slower release rather than the final amount released. Because we used five or more experimental replicates with multiple data points for each condition to calculate mean release rates, we estimated release error using the standard error of the mean (s.e.m.), whereas we used standard deviation (s.d.) to estimate error for measurements with only three replicates and data points in other assays. For the release measurements under oxidizing conditions, we calculated the release efficiencies of the crosslinked fractions of Cys-pair RNAP using the following equation:

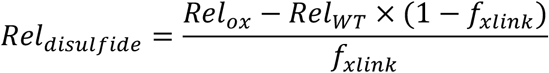

where *Rel*_*disulfide*_ is the corrected release efficiency of the Cys-pair crosslinked EC, *Rel*_*ox*_ is the observed release efficiency of the Cys-pair EC under oxidizing conditions (crosslinked + noncrosslinked), *Rel*_*WT*_ is the observed release efficiency of the WT EC under oxidizing conditions, and *f*_*xlink*_ is the crosslinking efficiency.

### MccJ25 abortive initiation assays

*In vitro* abortive initiation transcription assays were performed by incubating 150 nM wild-type RNAP σ^70^ holoenzyme with 75 nM *lac*UV5 promoter-containing DNA template in Cl^−^ elongation buffer at 37 °C for 15 min to form 75 nM RP_c_. MccJ25 was titrated into aliquots followed by incubation for an additional 5 min at 37 °C. Finally, an NTP mix was added to each aliquot for a final concentration of 25 nM RP_ITC_, 0–128 μM MccJ25, 500 μM ApA (TriLink BioTechnologies), 50 μM UTP, and 0.5 mCi [α-^32^P] UTP/μl. Reactions were stopped after 10 min by addition of an equal volume of 2X urea stop buffer, separated by electrophoresis in a denaturing 23% (v/v) polyacrylamide gel, and quantitated using a *via* PhosphorImager. Abortive RNAs were mapped as in Borowiec and Galla [61].

### Trigger loop cysteine-pair reporter assays

MccJ25 (100 μM) was added to 50 nM RNAP and incubated for 5 min at 37 °C. Disulfide formation was initiated by treatment with 0.25 mM DTT and 10 mM CSSC (*E*_h_ = −0.16 mV) for 60 min at 37 °C. Reactions were quenched with 50 mM iodoacetamide and separated by non-reducing SDS-PAGE using 8% Bolt Bis-Tris Plus gels. Gels were stained with Krypton Fluorescent Protein Stain and imaged with a 520-nm excitation filter and 580-nm emission filter. Crosslinking efficiency was determined using ImageQuant software. Experimental error was determined as the standard deviation of measurements from three independent replicates.

### Determination of commitment rates and propagation of error

TE was calculated as in Reynolds and Chamberlin [62] and redefined in terms of rates as in von Hippel and Yager [63] such that:

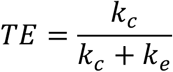

and

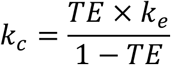

where *k*_c_ represents the termination commitment rate and *k*_e_ represents the elongation rate. We express elongation rate measurements here as relative pseudo-first-order rate constants, *k*_e_, for reactions that may encompass more than one step or for which concentrations are assumed to be unchanging and thus not represented. The relative pseudo-first-order rate constants will be proportional to composite rate constants for the reactions and are specific to our experimental conditions.

To determine *k*_c_, we first extracted overall elongation rates using KinTek Explorer. We calculated overall elongation rates by fitting the appearance of runoff transcript to the one-step, pseudo-first-order kinetic model of A26 → RO initially seeded with rates for A26 disappearance and RO appearance determined using for single-exponential fits using IgorPro (WaveMetrics). All replicates for a given RNAP variant and redox condition were fit simultaneously to minimize the experimental error. These average elongation rates were then normalized such that under reducing conditions the WT RNAP had the arbitrary *k*_e_ = 1 (see Figure 3B). We then determined *k*_c_ for each RNAP under both reducing and oxidizing conditions.

Because TE and *k*_e_ were separate measurements with separate associated errors, determination of *k*_c_ required propagation of uncertainty using the following equation:

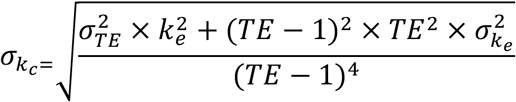

## ACKNOWLEDGEMENTS

We thank members of the Landick Group and the Record Group for helpful discussions and comments on the manuscript. M.J.B. was supported by the Molecular Biosciences Predoctoral Training Grant from the NIH (T32GM007215). This research was supported by a grant from the NIGMS to R.L. (GM38660).

---

**Figure S1.**
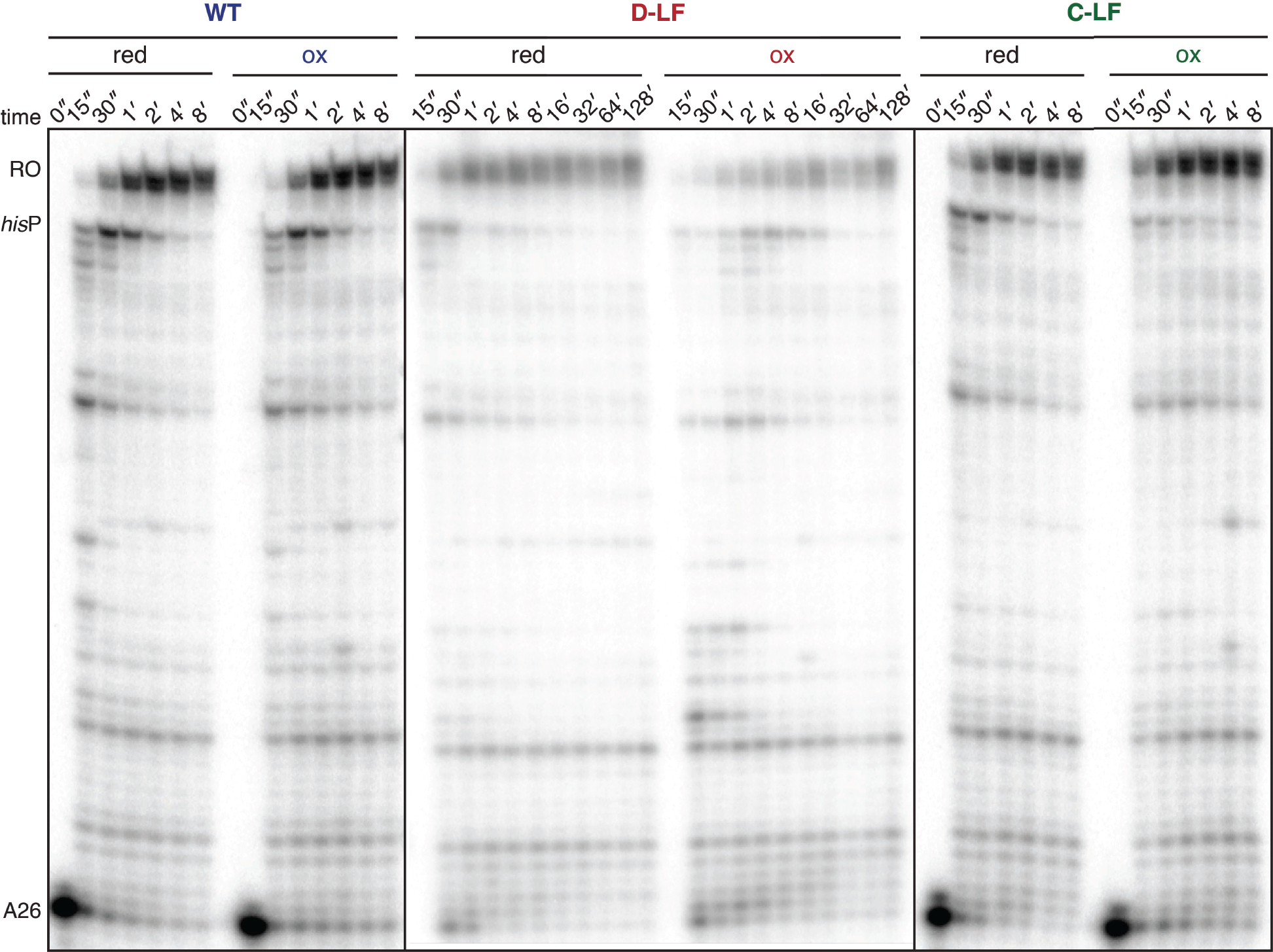
Constraining the clamp perturbs overall elongation rates through effects on relative pause strength. Representative gels showing the transcriptional behavior of WT, D-LF, and C-LF ECs under reducing and oxidizing conditions. RO, template run-off.

**Figure S2.**
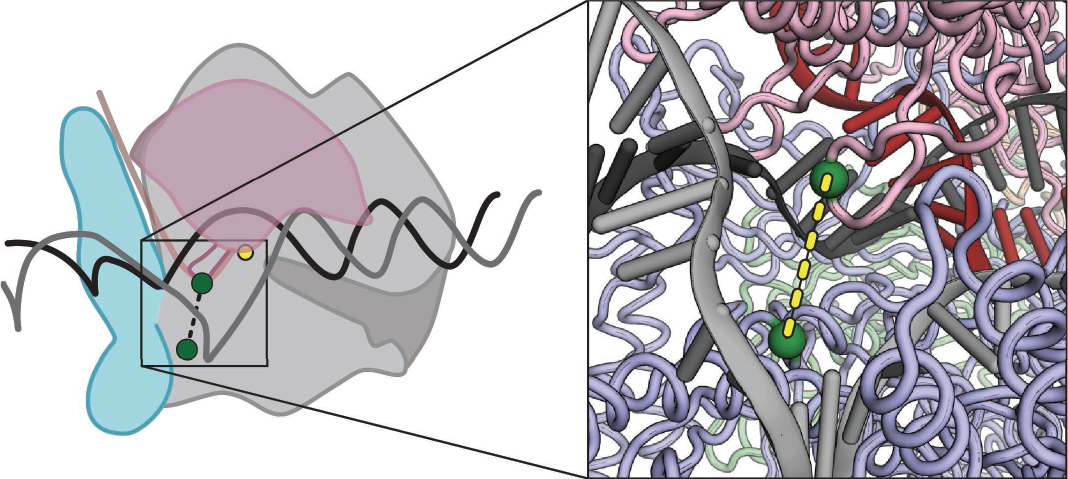
T-RP forms a disulfide crosslink within the transcription bubble. Left panel; position of the cysteine substitutions within the context of the entire EC. Right panel: zoomed-in view of the rudder-protrusion crosslink. Cα atoms are depicted as green spheres. A full nucleic-acid scaffold is modeled in based upon previous structural predictions [65] (RNA, red; template DNA, black; nontemplate, gray).

**Figure S3.**
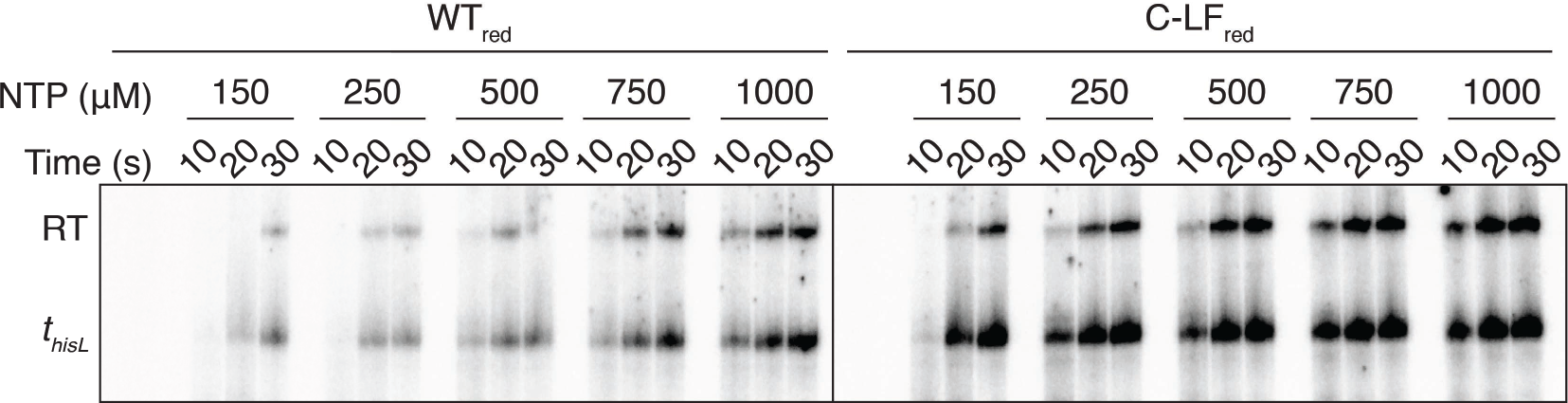
Elongation to *t*_*hisL*_ from A26 with different NTP concentrations under reducing conditions.

**Figure S4.**
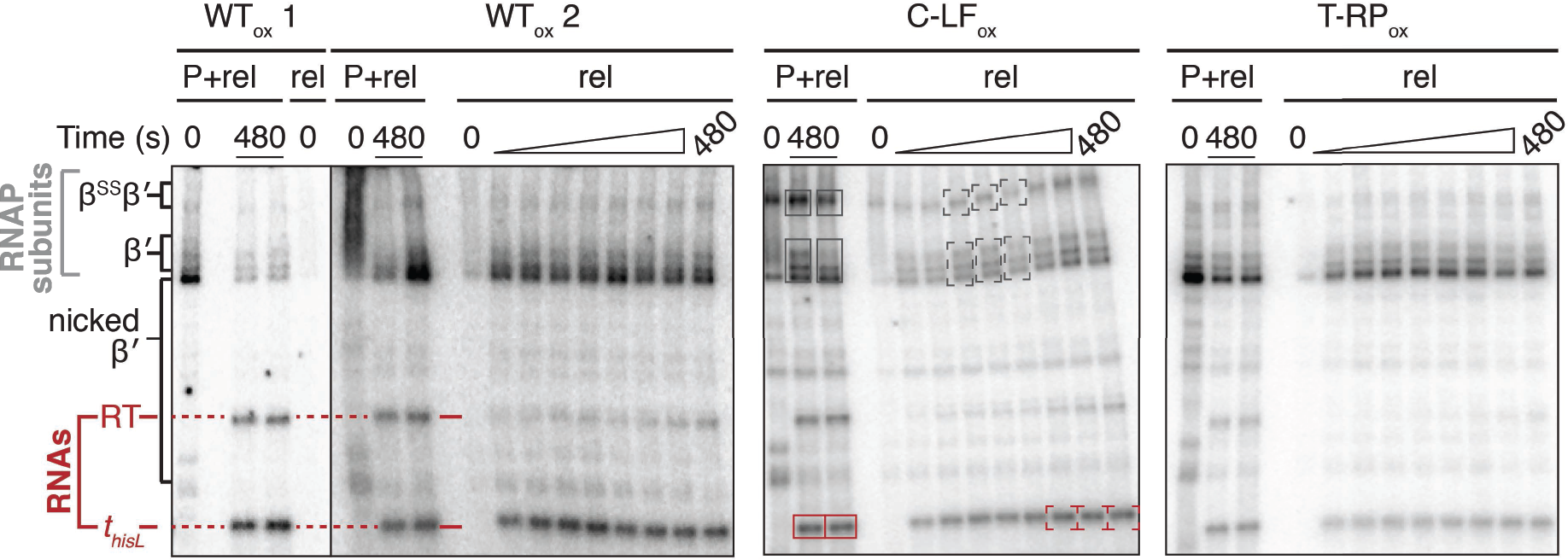
Representative dissociation assay non-reducing gels. P, bead-bound pellet; rel, sample <u>rel</u>eased into the reaction supernatant. Two sets of P+rel (total sample) and 0 s timepoints of rel samples for WT ECs are shown on the left (WT_ox_=1 and WT_ox_=2) with replicate P+rel samples from the 480 s timepoint in adjacent lanes. As seen in WT_ox_=2, aggregation sometimes complicated measurements of total sample RNAP amounts. RNAs used for quantitation are marked in red: *t*_hisL_, terminated RNA; RT, readthough RNA formed when ECs reach the end of the template. Proteolytic nicking of β′ generated minor and variable amounts of C-terminal ^32^P-labeled bands (nicked β′); these bands were not used for quantitiation. Because quantitation relied on comparison of the amounts of β′ or RNA in the total vs. supernatant fractions, the small amount of nicked β′ did not interfere with quantitation. Multiple full-length β′ bands sometimes appeared because the gels were not run under reducing conditions to allow detection of the crosslinked β′. We included these variant β′ bands, which likely reflect internal disulfide formation, when quantifying released RNAP. Release timepoints taken at 0, 20, 30, 40, 50, 60, 120, 240, and 480 s. The C-LF gel (middle panel) is marked with examples of the bands used to quantify RNA and RNAP release efficiencies. Red boxes and brackets indicate RNA species at *t*_*hisL*_ dark gray boxes and brackets indicate the ^32^P-β′ and ^32^P-β^SS^β′ bands. For this example, RNA release efficiency was calculated as the ^32^P signal minus background in red brackets divided by the average ^32^P signal minus background in the two red boxes. RNAP release efficiency is calculated as the ^32^P signal minus background in dark gray brackets divided by the ^32^P signal minus background in the dark gray boxes, multiplied by the termination efficiency to account for ECs that read through the terminator signal and did not release from the beads.

**Figure S5.**
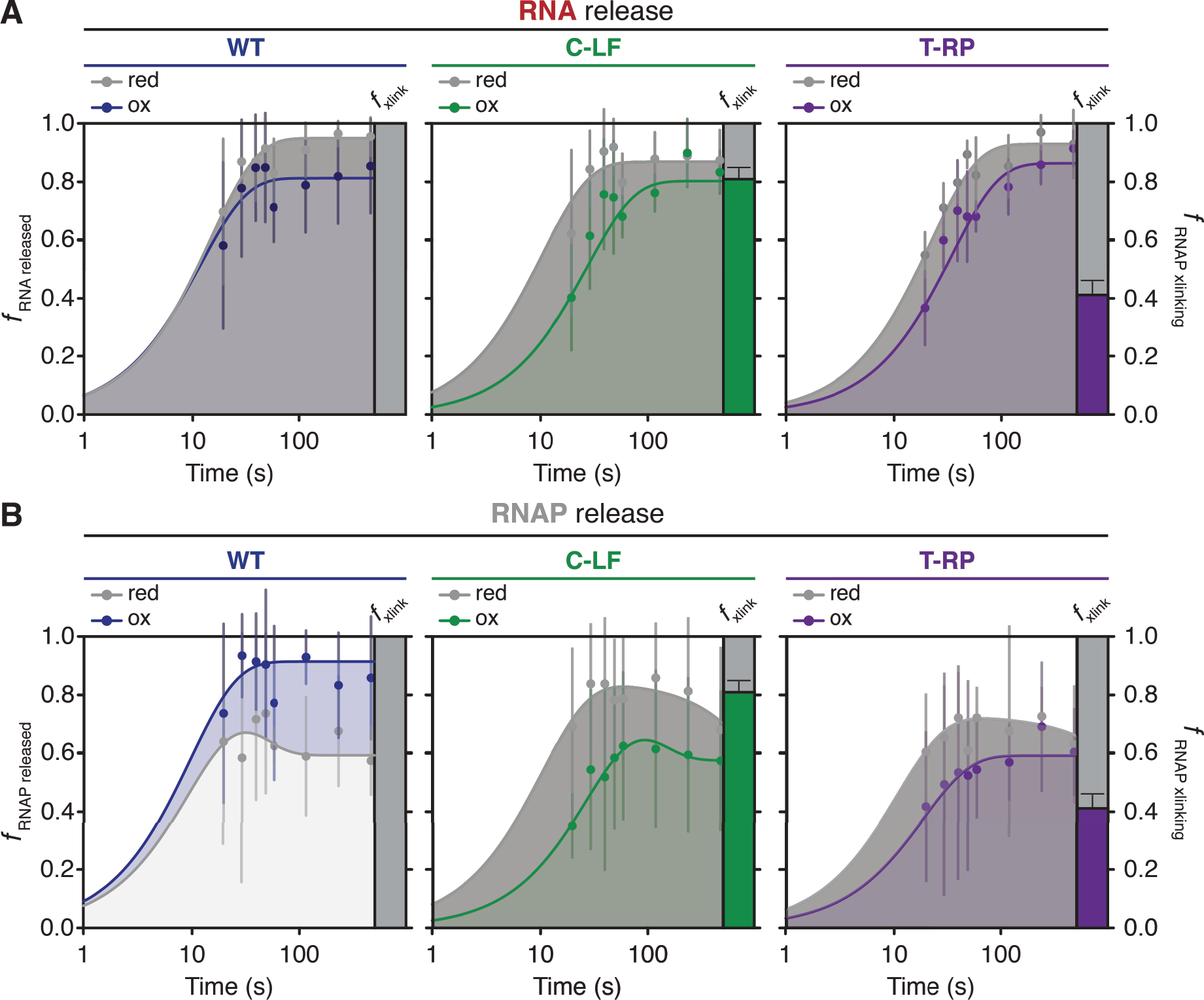
Evaluation of RNA and RNAP dissociation rates. (**A**) Semi-log plots of RNA release rates at *t*_*hisL*_. RNA dissociation was determined as the fraction of RNA released into the reaction supernatant relative to total RNA at the terminator site. Data were fit to either a single exponential [y = A(1 - e^−kt^)] function. The y-axis label “efficiency” represents measurements of both Cys-pair crosslinking efficiency and RNA release efficiency. Bars to the right of plots represent crosslinking efficiencies under oxidizing conditions. Grey curves represent fits for RNA release under reducing conditions; colored curves represent fits under oxidizing conditions. Error bars represent the standard deviation at n = 3. (**B**) Semi-log plots of RNAP release at *t*_*hisL*_. RNAP dissociation was determined as the fraction of RNAP released into the reaction supernatant relative to total RNAP within the reaction mixture multiplied by the TE. Data were fit to the double exponential function 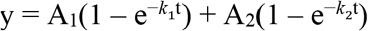. Plot components are the same as in subpanel A.

**Figure S6.**
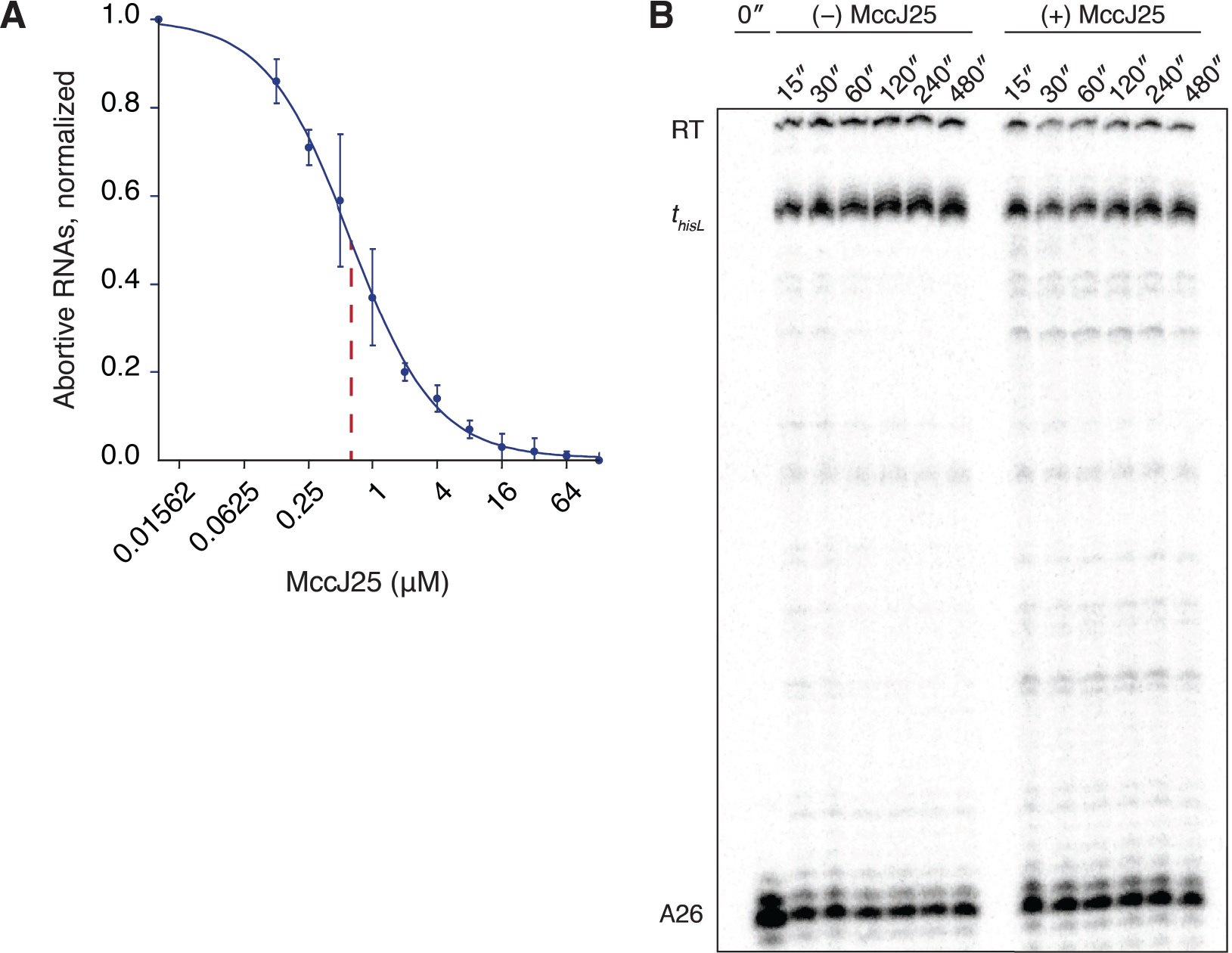
Effects of MccJ25 on RNAP during abortive initiation and elongation. (**A**) Abortive initiation by WT RNAP at *lac*UV5 is inhibited by MccJ25. Abortive products were normalized to the maximum signal in the absence of MccJ25. (**B**) Representative gel showing that the addition of MccJ25 to *in vitro* transcription assays at 5 s is sufficient time to allow elongation by the majority of ECs to *t*_*hisL*_ under conditions of saturating NTP concentration. The presence and persistence of pause bands in the +MccJ25 samples indicates successful arrest of lagging ECs by the inhibitor.

**Figure S7.**
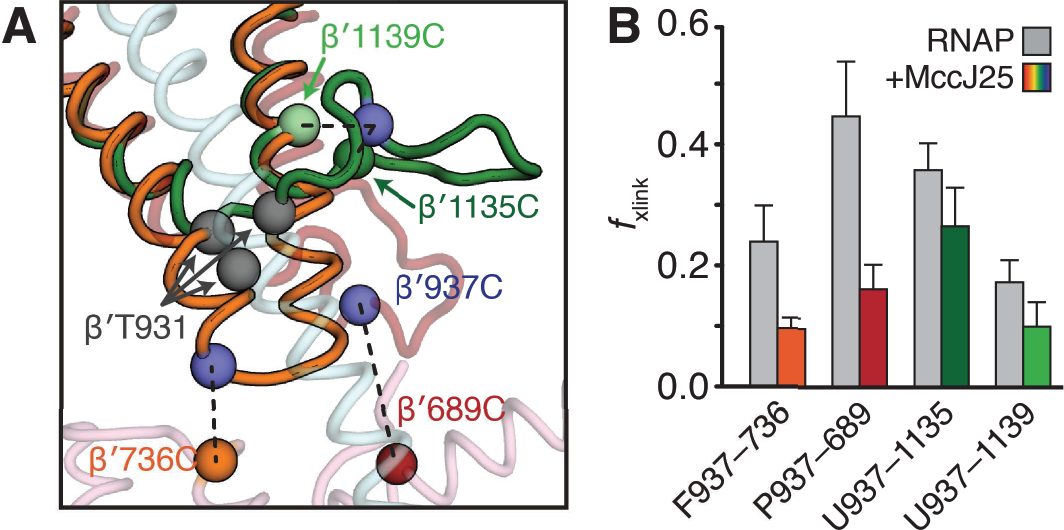
Cysteine-pair reporting suggests MccJ25 binds the TL in its unfolded conformation. (**A**) Location of TL cysteine-pairs. Three TL conformations observed in crystal structures were aligned to a cryo-EM structure of the *E. coli* EC [27]: orange, folded trigger helices, PDB ID: **<u>2Q5J</u>** [8]; red, partially-folded TL, PDB ID: **<u>2NVO</u>** [66]; green, unfolded TL, PDB ID: **<u>1IW7</u>** [38]. The bridge helix is depicted in cyan. Spheres depict the Cα atoms of relevant β′ residues. Grey spheres indicate the position of β′T931, a residue that when mutated (T931I) confers full resistance to MccJ25 and is thus likely a critical MccJ25 binding determinant. (**B**) Crosslinking efficiencies of cysteine-pair reporters in the absence (grey bars) and presence (colored bars) of MccJ25. Error represents the standard deviation (n=3).

## Abbreviations

EC: elongating transcription complex
*f*_xlink_: crosslinking efficiency
Cys-pair: cysteine-pair
MccJ25: microcin J25
NA: nucleic acid
RNAP: RNA polymerase
TE: termination (commitment) efficiency
T_hp_: terminator hairpin
TL: trigger loop

## REFERENCES

[1] Peters JM, Mooney RA, Grass JA, Jessen ED, Tran F, Landick R. Rho and NusG suppress pervasive antisense transcription in Escherichia coli. Genes & development. 2012;26:2621–33.

[2] Helmrich A, Ballarino M, Nudler E, Tora L. Transcription-replication encounters, consequences and genomic instability. Nature structural & molecular biology. 2013;20:412–8.

[3] d’Aubenton Carafa Y, Brody E, Thermes C. Prediction of rho-independent Escherichia coli transcription terminators. A statistical analysis of their RNA stem-loop structures. J Mol Biol. 1990;216:835–58.

[4] Wilson KS, von Hippel PH. Transcription termination at intrinsic terminators: the role of the RNA hairpin. Proc Natl Acad Sci U S A. 1995;92:8793–7.

[5] Yager TD, von Hippel PH. A thermodynamic analysis of RNA transcript elongation and termination in Escherichia coli. Biochemistry. 1991;30:1097–118.

[6] Komissarova N, Becker J, Solter S, Kireeva M, Kashlev M. Shortening of RNA:DNA hybrid in the elongation complex of RNA polymerase is a prerequisite for transcription termination. Molecular cell. 2002;10:1151–62.

[7] Korzheva N, Mustaev A, Kozlov M, Malhotra A, Nikiforov V, Goldfarb A, et al. A structural model of transcription elongation. Science. 2000;289:619–25.

[8] Vassylyev D, Vassylyeva M, Zhang J, Palangat M, Artsimovitch I, Landick R. Structural basis for substrate loading in bacterial RNA polymerase Nature 2007;448:163–8.

[9] Nudler E, Avetissova E, Markovtsov V, Goldfarb A. Transcription processivity: protein-DNA interactions holding together the elongation complex. Science. 1996;273:211–7.

[10] Gusarov I, Nudler E. The mechanism of intrinsic transcription termination. Molecular cell. 1999;3:495–504.

[11] Yarnell WS, Roberts JW. Mechanism of intrinsic transcription termination and antitermination. Science. 1999;284:611–5.

[12] Larson MH, Mooney RA, Peters JM, Windgassen T, Nayak D, Gross CA, et al. A pause sequence enriched at translation start sites drives transcription dynamics in vivo. Science. 2014;344:1042–7.

[13] Toulokhonov I, Zhang J, Palangat M, Landick R. A central role of the RNA polymerase trigger loop in active-site rearrangement during transcriptional pausing. Molecular cell. 2007;27:406–19.

[14] Ray-Soni A, Mooney RA, Landick R. Trigger loop dynamics can explain stimulation of intrinsic termination by bacterial RNA polymerase without terminator hairpin contact. Proc Natl Acad Sci U S A. 2017;114:E9233–E42.

[15] Martin FH, Tinoco I, Jr. DNA-RNA hybrid duplexes containing oligo(dA:rU) sequences are exceptionally unstable and may facilitate termination of transcription. Nucleic Acids Res. 1980;8:2295–9.

[16] Larson MH, Greenleaf WJ, Landick R, Block SM. Applied force reveals mechanistic and energetic details of transcription termination. Cell. 2008;132:971–82.

[17] Kashlev M, Komissarova N. Transcription termination: primary intermediates and secondary adducts. J Biol Chem. 2002;277:14501–8.

[18] Santangelo TJ, Roberts JW. Forward translocation is the natural pathway of RNA release at an intrinsic terminator. Molecular cell. 2004;14:117–26.

[19] Epshtein V, Cardinale CJ, Ruckenstein AE, Borukhov S, Nudler E. An allosteric path to transcription termination. Molecular cell. 2007;28:991–1001.

[20] Epshtein V, Dutta D, Wade J, Nudler E. An allosteric mechanism of Rho-dependent transcription termination. Nature. 2010;463:245–9.

[21] Weixlbaumer A, Leon K, Landick R, Darst SA. Structural basis of transcriptional pausing in bacteria. Cell. 2013;152:431–41.

[22] Chakraborty A, Wang D, Ebright YW, Korlann Y, Kortkhonjia E, Kim T, et al. Opening and closing of the bacterial RNA polymerase clamp. Science. 2012;337:591–5.

[23] Gnatt AL, Cramer P, Fu J, Bushnell DA, Kornberg RD. Structural basis of transcription: an RNA polymerase II elongation complex at 3.3 A resolution. Science. 2001;292:1876–82.

[24] Sekine S, Murayama Y, Svetlov V, Nudler E, Yokoyama S. The ratcheted and ratchetable structural states of RNA polymerase underlie multiple transcriptional functions. Molecular cell. 2015;57:408–21.

[25] Tagami S, Sekine S, Kumarevel T, Hino N, Murayama Y, Kamegamori S, et al. Crystal structure of bacterial RNA polymerase bound with a transcription inhibitor protein. Nature. 2010;468:978–82.

[26] Vassylyev DG, Vassylyeva MN, Perederina A, Tahirov TH, Artsimovitch I. Structural basis for transcription elongation by bacterial RNA polymerase. Nature. 2007;448:157–62.

[27] Kang JY, Olinares PD, Chen J, Campbell EA, Mustaev A, Chait BT, et al. Structural basis of transcription arrest by coliphage HK022 Nun in an Escherichia coli RNA polymerase elongation complex. Elife. 2017;6.

[28] Weilbaecher R, Hebron C, Feng G, Landick R. Termination-altering amino acid substitutions in the beta’ subunit of Escherichia coli RNA polymerase identify regions involved in RNA chain elongation. Genes & development. 1994;8:2913–27.

[29] Cheeran A, Babu Suganthan R, Swapna G, Bandey I, Achary MS, Nagarajaram HA, et al. Escherichia coli RNA polymerase mutations located near the upstream edge of an RNA:DNA hybrid and the beginning of the RNA-exit channel are defective for transcription antitermination by the N protein from lambdoid phage H-19B. J Mol Biol. 2005;352:28–43.

[30] Kang JY, Mishanina TV, Bellecourt MJ, Mooney RA, Darst SA, Landick R. RNA Polymerase Accommodates a Pause RNA Hairpin by Global Conformational Rearrangements that Prolong Pausing. Molecular cell. 2018;69:802–15 e1.

[31] Hein PP, Kolb KE, Windgassen T, Bellecourt MJ, Darst SA, Mooney RA, et al. RNA polymerase pausing and nascent-RNA structure formation are linked through clamp-domain movement. Nature structural & molecular biology. 2014;21:794–802.

[32] Toulokhonov I, Landick R. The role of the lid element in transcription by E. coli RNA polymerase. J Mol Biol. 2006;361:644–58.

[33] Guo X, Myasnikov AG, Chen J, Crucifix C, Papai G, Takacs M, et al. Structural Basis for NusA Stabilized Transcriptional Pausing. Molecular cell. 2018;69:816–27 e4.

[34] Palangat M, Larson MH, Hu X, Gnatt A, Block SM, Landick R. Efficient reconstitution of transcription elongation complexes for single-molecule studies of eukaryotic RNA polymerase II. Transcription. 2012;3:146–53.

[35] Mishanina TV, Palo MZ, Nayak D, Mooney RA, Landick R. Trigger loop of RNA polymerase is a positional, not acid-base, catalyst for both transcription and proofreading. Proc Natl Acad Sci U S A. 2017;114:E5103–E12.

[36] Nayak D, Voss M, Windgassen T, Mooney RA, Landick R. Cys-pair reporters detect a constrained trigger loop in a paused RNA polymerase. Molecular cell. 2013;50:882–93.

[37] Cramer P, Bushnell D, Kornberg R. Structural basis of transcription: RNA polymerase II at 2.8 Å resolution. Science. 2001;292:1863–76.

[38] Vassylyev DG, Sekine S, Laptenko O, Lee J, Vassylyeva MN, Borukhov S, et al. Crystal structure of a bacterial RNA polymerase holoenzyme at 2.6 A resolution. Nature. 2002;417:712’9.

[39] Felklistov A, Bae B, Hauver J, Lass-Napiorkowska A, Kalesse M, Glaus F, et al. RNA polymerase motions during promoter melting. Science. 2017;356:863–6.

[40] Bayro MJ, Mukhopadhyay J, Swapna G, Huang JY, Ma L, Sineva E, et al. Structure of antibacterial peptide microcin J25: a 21-residue lariat protoknot. J Am Chem Soc. 2003;125:12382–3.

[41] Mukhopadhyay J, Sineva E, Knight J, Levy RM, Ebright RH. Antibacterial peptide microcin J25 inhibits transcription by binding within and obstructing the RNA polymerase secondary channel. Molecular cell. 2004;14:739–51.

[42] Yuzenkova J, Delgado M, Nechaev S, Savalia D, Epshtein V, Artsimovitch I, et al. Mutations of bacterial RNA polymerase leading to resistance to microcin j25. J Biol Chem. 2002;277:50867–75.

[43] Delgado MA, Rintoul MR, Farias RN, Salomon RA. Escherichia coli RNA polymerase is the target of the cyclopeptide antibiotic microcin J25. J Bacteriol. 2001;183:4543–50.

[44] Windgassen TA, Mooney RA, Nayak D, Palangat M, Zhang J, Landick R. Trigger-helix folding pathway and SI3 mediate catalysis and hairpin-stabilized pausing by Escherichia coli RNA polymerase. Nucleic Acids Res. 2014;42:12707–21.

[45] Kang JY, Mooney RA, Nedialkov Y, Saba J, Mishanina TV, Artsimovitch I, et al. Structural Basis for Transcript Elongation Control by NusG Family Universal Regulators. Cell. 2018;173:1650–62.e14.

[46] Shetty A, Kallgren SP, Demel C, Maier KC, Spatt D, Alver BH, et al. Spt5 Plays Vital Roles in the Control of Sense and Antisense Transcription Elongation. Molecular cell. 2017;66:77–88 e5.

[47] Yamaguchi Y, Mura T, Chanarat S, Okamoto S, Handa H. Hepatitis delta antigen binds to the clamp of RNA polymerase II and affects transcriptional fidelity. Genes Cells. 2007;12:863–75.

[48] Singh N, Bubunenko M, Smith C, Abbott DM, Stringer AM, Shi R, et al. SuhB Associates with Nus Factors To Facilitate 30S Ribosome Biogenesis in Escherichia coli. MBio. 2016;7:e00114.

[49] Arnvig KB, Zeng S, Quan S, Papageorge A, Zhang N, Villapakkam AC, et al. Evolutionary comparison of ribosomal operon antitermination function. J Bacteriol. 2008;190:7251–7.

[50] Peters JM, Vangeloff AD, Landick R. Bacterial transcription terminators: the RNA 3’-end chronicles. J Mol Biol. 2011;412:793–813.

[51] Kireeva M, Trang C, Matevosyan G, Turek-Herman J, Chasov V, Lubkowska L, et al. RNA-DNA and DNA-DNA base-pairing at the upstream edge of the transcription bubble regulate translocation of RNA polymerase and transcription rate. Nucleic Acids Res. 2018;46:5764–75.

[52] Ryder AM, Roberts JW. Role of the Non-template Strand of the Elongation Bubble in Intrinsic Transcription Termination. Journal of Molecular Biology. 2003;334:205–13.

[53] Harden TT, Wells CD, Friedman LJ, Landick R, Hochschild A, Kondev J, et al. Bacterial RNA polymerase can retain sigma70 throughout transcription. Proc Natl Acad Sci U S A. 2016;113:602–7.

[54] Deighan P, Pukhrambam C, Nickels BE, Hochschild A. Initial transcribed region sequences influence the composition and functional properties of the bacterial elongation complex. Genes & development. 2011;25:77–88.

[55] Bar-Nahum G, Nudler E. Isolation and Characterization of σ70-Retaining Transcription Elongation Complexes from Escherichia coli. Cell. 2001;106:443–51.

[56] Malinen AM, Nandymazumdar M, Turtola M, Malmi H, Grocholski T, Artsimovitch I, et al. CBR antimicrobials alter coupling between the bridge helix and the beta subunit in RNA polymerase. Nat Commun. 2014;5:3408.

[57] Gribskov M, Burgess RR. Overexpression and purification of the sigma subunit of Eschevichia coli RNA polymerase. Gene. 1985;26:109–18.

[58] Pan SJ, Cheung WL, Link AJ. Engineered gene clusters for the production of the antimicrobial peptide microcin J25. Protein Expr Purif. 2010;71:200–6.

[59] Hoopes BC, McClure WR. Studies on the selectivity of DNA precipitation by spermine. Nucleic Acids Res. 1981;9:5493–504.

[60] Borukhov S, Laptenko O, Lee J. Escherichia coli Transcript Cleavage Factors GreA and GreB: Functions and Mechanisms of Action. Ribonucleases - Part B 2001. p. 64–76.

[61] Borowiec JA, Gralla JD. Supercoiling response of the lac ps promoter in vitro. J Mol Biol. 1985;184:597–8.

[62] Reynolds R, Bermudez-Cruz RM, Chamberlin MJ. Parameters affecting transcription termination by Escherichia coli RNA polymerase. I. Analysis of 13 rho-independent terminators. J Mol Biol. 1992;224:31–51.

[63] von Hippel PH, Yager TD. Transcript elongation and termination are competitive kinetic processes. Proc Natl Acad Sci U S A. 1991;88:2307–11.

[64] Ederth J, Artsimovitch I, Isaksson LA, Landick R. The downstream DNA jaw of bacterial RNA polymerase facilitates both transcriptional initiation and pausing. J Biol Chem. 2002;277:37456–63.

[65] Opalka N, Brown J, Lane WJ, Twist KA, Landick R, Asturias FJ, et al. Complete structural model of Escherichia coli RNA polymerase from a hybrid approach. PLoS Biol. 2010;8.

[66] Wang D, Bushnell DA, Westover KD, Kaplan CD, Kornberg RD. Structural basis of transcription: role of the trigger loop in substrate specificity and catalysis. Cell. 2006;127:941–54.

